# Beat encoding at mistuned octaves within single sensory neurons

**DOI:** 10.1101/2022.08.18.504442

**Authors:** Alexandra Barayeu, Ramona Schäfer, Jan Grewe, Jan Benda

## Abstract

Beats are periodic amplitude modulations resulting from the superposition of two spectrally close periodic signals, where the difference between the two signal frequencies defines the frequency of the beat. Such amplitude modulations are known to be well encoded in corresponding firing rate modulations in auditory fibers as well as in electrosensory afferents. In a field study we showed the behavioral relevance of beat-like amplitude modulations exceeding spectrally close interactions in electric fish. Thus, we here study the encoding of beat-like waveforms over a wide range of difference frequencies in the electrosensory system of *Apteronotus leptorhynchus*. Contrary to expectations from the previously measured beat tuning, the activity of p-type electroreceptor afferents follows a repetitive pattern with slow modulations of their firing rate reoccurring around multiples of the frequency of the carrier signal. Mathematical reasoning supported by simulations of modified integrate-and-fire models reveals that neither Hilbert transform, squaring, harmonics of the carrier, half-wave rectification, nor the threshold-non-linearity of a spike generator are sufficient to extract slow beating signal envelopes around the octave of the carrier. Rather, a threshold operation smoothed out by exponentiation with a power of three is needed prior to spike generation to explain the repetitive occurrence of slow signal envelopes and electroreceptor responses. Our insights suggest the synapses of inner hair cells as candidate mechanisms underlying the perception of beats at mistuned octaves that has been described by Georg Simon Ohm, Hermann Helmholtz, and others already in the 19th century.

## 1. Introduction

Periodic signals are ubiquitous in the auditory system (Lewicki, 2002; Köoppl, 1997; Romani et al., 1982) and the electrosensory system of wave-type electric fish (Hopkins, 1976). The superposition of two periodic signals with similar frequencies results in a periodic amplitude modulation (AM) known as “beat”. The frequency of such a beat is given by the frequency difference between the two signals and the beat amplitude is the one of the smaller of the two signals. Auditory beats give rise to a unique beating perception (Roeber, 1834; Plomp, 1967). In wave-type electric fish, beats play a central role in electrocommunication (Benda, 2020).

Wave-type Gymnotiform electric fish generate a sinusoidal electric organ discharge (EOD) of a species and individual specific frequency (Knudsen, 1975; Henninger et al., 2020). The EODs of two nearby fish superimpose and thus produce a beat. Beat amplitude declines with distance between the two fish and thus is strongly influenced by relative movement (Yu et al., 2005). The periodic beat is modulated by various types of electrocommunication signals on time scales ranging from 10 ms to many seconds (Zakon et al., 2002; Smith, 2013). Cutaneous tuberous organs that are distributed all over the body (Carr et al., 1982) sense the actively generated electric field and its modulations. Within a single tuberous electroreceptor organ about 30 primary electroreceptors form ribbon synapses onto the dendrites of the same afferent fiber (Szabo, 1965; Wachtel and Szamier, 1966) which projects via the lateral line nerve to the hindbrain. There it synapses onto pyramidal neurons in the electrosensory lateral line lobe (Maler, 2009). So far, the locus of action potential generation is not known, but most likely right after where the dendrites merge into a single fiber.

The time course of the activity of the p-type electroreceptor afferents (P-units) follows the time course of AMs of the EOD (Bastian, 1981; Nelson et al., 1997; Benda et al., 2005), similar to auditory fibers (Joris et al., 2004). So far, P-unit tuning to beat frequencies has been analyzed in a range up to 300 Hz in *Apteronotus leptorhynchus*. Beyond the strongest firing rate modulations in response to beat frequencies of 60 – 100 Hz the response declines down to baseline at about 250 Hz (Bastian, 1981; Benda et al., 2006; Walz et al., 2014). Recent field studies on *Apteronotus*, however, demonstrated behaviorally relevant difference frequencies beyond 300 Hz in the context of courtship and synchronization of spawning (Henninger et al., 2018) and potential inter-species detection (Henninger et al., 2020).

Here we address the apparent mismatch between P-unit tuning and behavioral relevant beat frequencies. We record P-unit activities of *Apteronotus lepthorynchus* in response to a much wider range of difference frequencies (−750 ≤ Δ*f* ≤ 2500 Hz) than before, also exceeding the ranges observed for inter- and intra species encounters in the field (Henninger et al., 2018, 2020). Slow, beat-like envelopes repetitively reoccur at multiples of the carrier EOD and P-unit firing follow this pattern up to at least three multiples of the EOD frequency. By mathematical reasoning and simulations of integrate-and-fire neurons (Chacron et al., 2001; Savard et al., 2011; Sinz et al., 2020) we identify the minimum functional requirements on single sensory cells to be able to extract and respond to the slow envelopes generated by large difference frequencies. Finally, we test our insights in behavioral experiments based on the jamming avoidance response (Watanabe and Takeda, 1963).

Our results generate an interesting hypothesis for the mechanism underlying the perception of beats at mistuned octaves in humans. Experiments dating back to the 19th century demonstrated a perception of beats not only for spectrally close frequencies, but also for mistuned octaves where the second frequency is close to octaves of the carrier frequency (Roeber, 1834; König, 1876; Ohm, 1839; Helmholtz, 2009). The physiological mechanisms underlying these percepts remain an open issue (Plomp, 1967; Joris et al., 2004).

## 2. Results

We studied the encoding of beats over a wide range of frequencies by presenting sinusoidal electrical stimuli with absolute frequencies ranging from 20 up to 3200 Hz to electric fish of the species *A. leptorhynchus*. The stimuli superimpose with the fish’s own electric field and result in beat-like envelopes. In the following we speak of beats for difference frequencies below half an octave and of beat-like envelopes for larger difference frequencies. We recorded the spiking activity of *n* = 40 p-type electroreceptors afferents (P-units) in response to these stimuli.

### Responses to low difference frequencies

P-units are known to respond to low-frequency beats by modulating their firing rate (Bastian, 1981; Nelson et al., 1997; Benda et al., 2005; Walz et al., 2014). For a fish with an EOD frequency *f_EOD_* = 668 Hz and a stimulus with frequency *f_stim_* = 737 Hz, mimicking a fish with higher EOD frequency, we get a beat at the difference frequency Δ*f* = *f_stim_ − f_EOD_* = 69 Hz (Fig. 1 C, top). A P-unit responds to this amplitude modulation as demonstrated by the spike raster and the corresponding time-resolved firing rate (Fig. 1 C, center). The strongest peaks in the power spectrum of the spike response are at the beat frequency (colored marker) and the receiver’s EOD frequency (gray circle, Fig. 1 C bottom).

**Figure 1:**
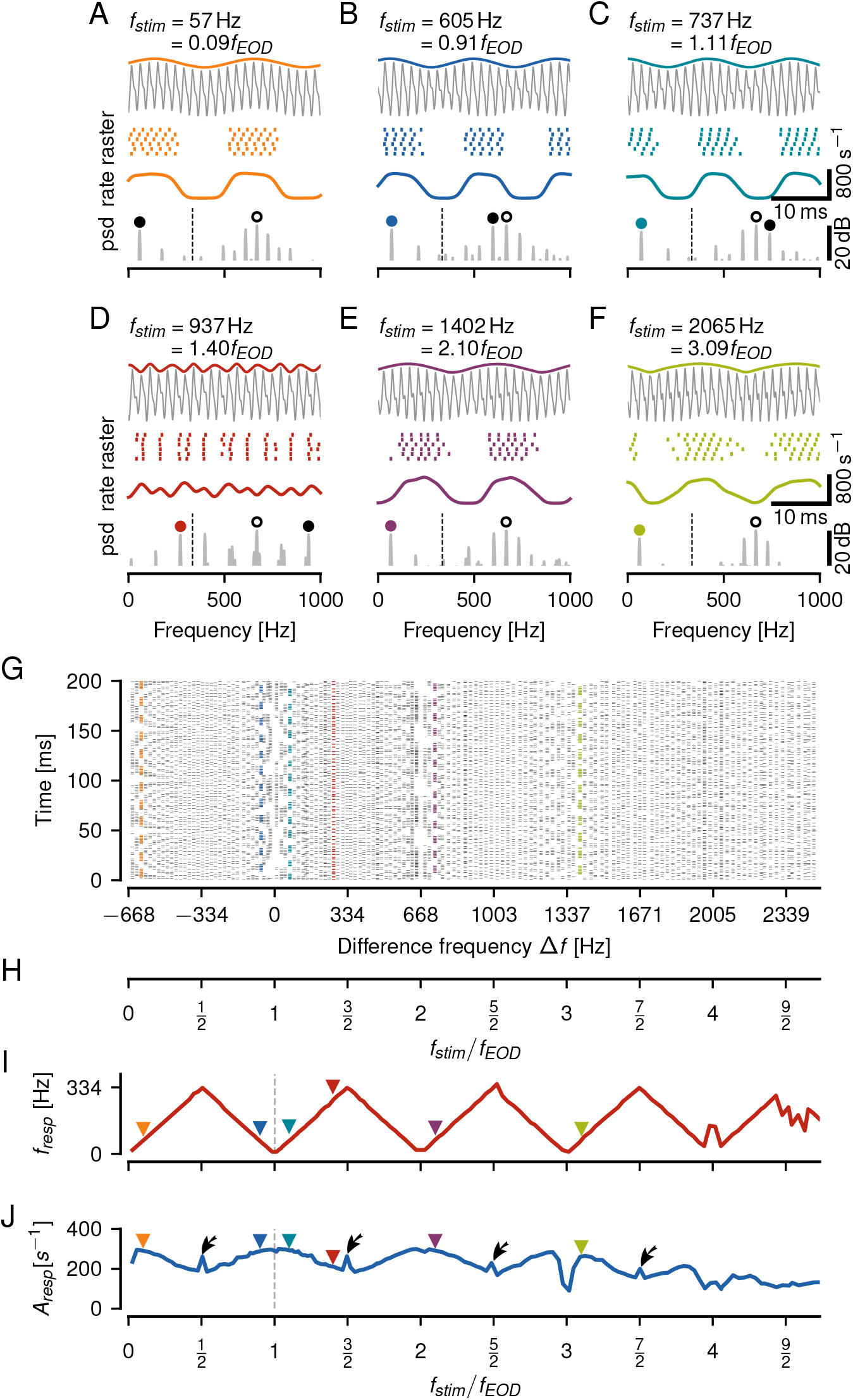
Encoding of a wide range of difference frequencies in an example P-unit. **A–F** Beating envelopes (colored lines) of the carrier EOD superimposed with a sine wave stimulus (gray, top), spike raster and firing rate of evoked P-unit responses (center), and corresponding power spectrum (psd, bottom, open circle: *f_EOD_*, dashed line: *f_EOD_*/2, colored circle: strongest frequency below *f_EOD_*/2) of a few selected stimulus frequencies as indicated. **G** Vertical raster plots showing spiking responses to a wide range of stimulus frequencies (one trial per frequency) indicate a repetitive structure of P-unit responses. Colored rasters mark examples shown in panels A–F. **H** Alternative stimulus frequency axis in multiples of *f_EOD_*, i.e. Δ*f/f_EOD_* + 1. **I** Frequency tuning, *f_resp_*, of the P-unit response, i.e. frequency of its firing rate modulation, retrieved as the strongest peak in the response spectrum below *f_EOD_*/2, repeats every integer multiple of *f_EOD_*. Colored triangles mark examples shown in panels A–F. **J** Amplitude tuning curve, quantified as the amplitude of the peak at *f_resp_* (square-root of the integral below the peak at *f_resp_* of the power spectrum of the spike response convolved with a Gaussian kernel with *σ* = 0.5 ms), also repeats at harmonics of *f_EOD_*. Strongest responses are close to multiples of *f_EOD_*. Right at odd multiples of *f_EOD_*/2 responses are enhanced (arrows).

When *f_stim_* is below *f_EOD_*(here Δ*f* = −63 Hz), the resulting beat is similar to the one discussed above, although at a negative difference frequency. The P-unit response has the same features as for the positive difference frequency and the spectrum has prominent peaks at the same locations (compare Figs. 1 B and C). Signals generated by positive or negative Δ*f* differ only in small phase shifts in the carrier, encoded by another population of electroreceptors, the T-units (Scheich et al., 1973).

As *f_stim_* increases, the beat frequency increases accordingly and the strength of the P-unit’s response, characterized by the modulation depth of the P-unit’s firing rate, declines (Fig. 1 D), confirming previous results (Bastian, 1981; Nelson et al., 1997; Savard et al., 2011; Walz et al., 2014).

### Envelope frequency does not match difference frequency for high stimulus frequencies

In the examples discussed so far, |Δ*f*| and the frequency of the induced beating envelope are identical. Increasing (or decreasing) *f_stim_* beyond *f_EOD_* ± *f_EOD_*/2, however, breaks this relation. Instead, at *f_stim_* = 0.1*f_EOD_*, 2.1*f_EOD_*, or 3.1*f_EOD_* the resulting beating envelopes have the same frequency as for *f_stim_* = 1.1*f_EOD_*, the classical beat for a stimulus frequency close to the receiver’s EOD frequency (compare Fig. 1 A, E, F to C). Hence, even for extremely high difference frequencies the resulting envelopes can be rather slow.

### P-units respond to an extremely wide range of stimulus frequencies

Such slow envelopes effectively modulate the P-unit’s spike responses. This is well known for |Δ*f*| less than *f_EOD_*/2. Beyond such beats we observe reoccurring Δ*f* ranges that lead to strongly modulated responses up to difference frequencies of approximately 1400 Hz (Fig. 1 G). Also, towards large negative difference frequencies, corresponding to absolute stimulus frequencies down to about 35 Hz, we observed clear responses (Fig. 1 A, G). Δ*f* ranges leading to clear response modulations are centered on integer multiples of *f_EOD_*. Thus, *f_EOD_* defines the frequency scale in which *f_stim_* has to be interpreted. Accordingly, from now on we express *f_stim_* relative to *f_EOD_*, which also allows for comparisons across animals with distinct EOD frequencies (Fig. 1 H).

### Aliasing structure of beat responses

Around integer multiples of *f_EOD_* we see slow beat-like signals and correspondingly strongly modulated spike responses. Towards odd multiples of *f_EOD_*/2, signal envelopes get faster and spike responses get weaker. We quantified P-unit response characteristics by extracting the frequency and the respective strength of the response modulation from the power spectrum of the spiking response. The position of the strongest peak in the response power spectrum below *f_EOD_*/2 is the frequency of the P-unit’s firing modulation. Plotting this frequency as a function of *f_stim_* reveals a repetitive pattern shaped like a “Toblerone” (Fig. 1 I). This zigzag pattern is reminiscent of aliasing known from the sampling theorem, with *f_EOD_*/2 playing the role of the Nyquist frequency.

In a few cases the peak at the aliased frequency of the stimulus in a P-unit’s response spectrum is smaller than the peak at the baseline firing rate. This may happen at stimulus frequencies close to multiples of *f_EOD_* and this is why sometimes the frequency tuning curve deviates from the zigzag pattern (Fig. 1 I at 2005 Hz).

### Periodic amplitude tuning curve

The amplitude *A_resp_* of the respective peak in the response spectrum reflects the strength of the P-unit response, i.e. the modulation depth of its time-resolved firing rate. The resulting tuning curve also shows a repetitive structure (Fig. 1 J). Close to multiples of *f_EOD_* the response is strongest. These maxima, however, become smaller the higher *f_stim_*. Right at the multiples we observe dips in the response amplitudes which can be attributed to the P-unit’s spike-frequency adaptation (Benda et al., 2005). Response amplitudes decline as the stimulus frequency approaches odd multiples of *f_EOD_*/2. Exactly at odd multiples of *f_EOD_*/2, response amplitudes are often markedly elevated (arrows in Fig. 1 J). For higher stimulus frequencies response amplitude increases again towards the next multiple of *f_EOD_*. This increase of the response amplitude beyond 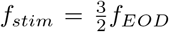 was not expected given previous data on the encoding of amplitude modulations in P-units (Bastian, 1981; Nelson et al., 1997; Savard et al., 2011; Walz et al., 2014).

### Amplitude tuning depends on post-synaptic filtering

The synapse between P-unit electroreceptor afferents and their target neurons in the hindbrain, pyramidal neurons in the electrosensory lateral line lobe, introduces a fast excitatory postsynaptic potential of about 1 ms duration (Berman and Maler, 1998). Postsynaptic potentials are the physiological equivalent of bins or filter kernels used to compute time-resolved firing rates. The shape of the P-unit’s amplitude tuning curve strongly depends on the width of the chosen filter kernel, because it low-pass filters the spike train, as does a postsynaptic potential. P-unit tuning is relatively flat when computed directly from the spike trains (no low-pass filtering, Fig. 2 A), but becomes more modulated the more the spike train is low-pass filtered, because the wider the postsynaptic potential, the more the response peaks closer to *f_EOD_*/2 are attenuated (arrows in Fig. 2 B,C). Low-pass filtering the spike train with a physiologically plausible Gaussian kernel with *σ* = 0.5 ms attenuates responses to high envelope frequencies while leaving responses to low envelope frequencies untouched, resulting in a periodically modulated amplitude tuning curve (Fig. 2 B).

**Figure 2:**
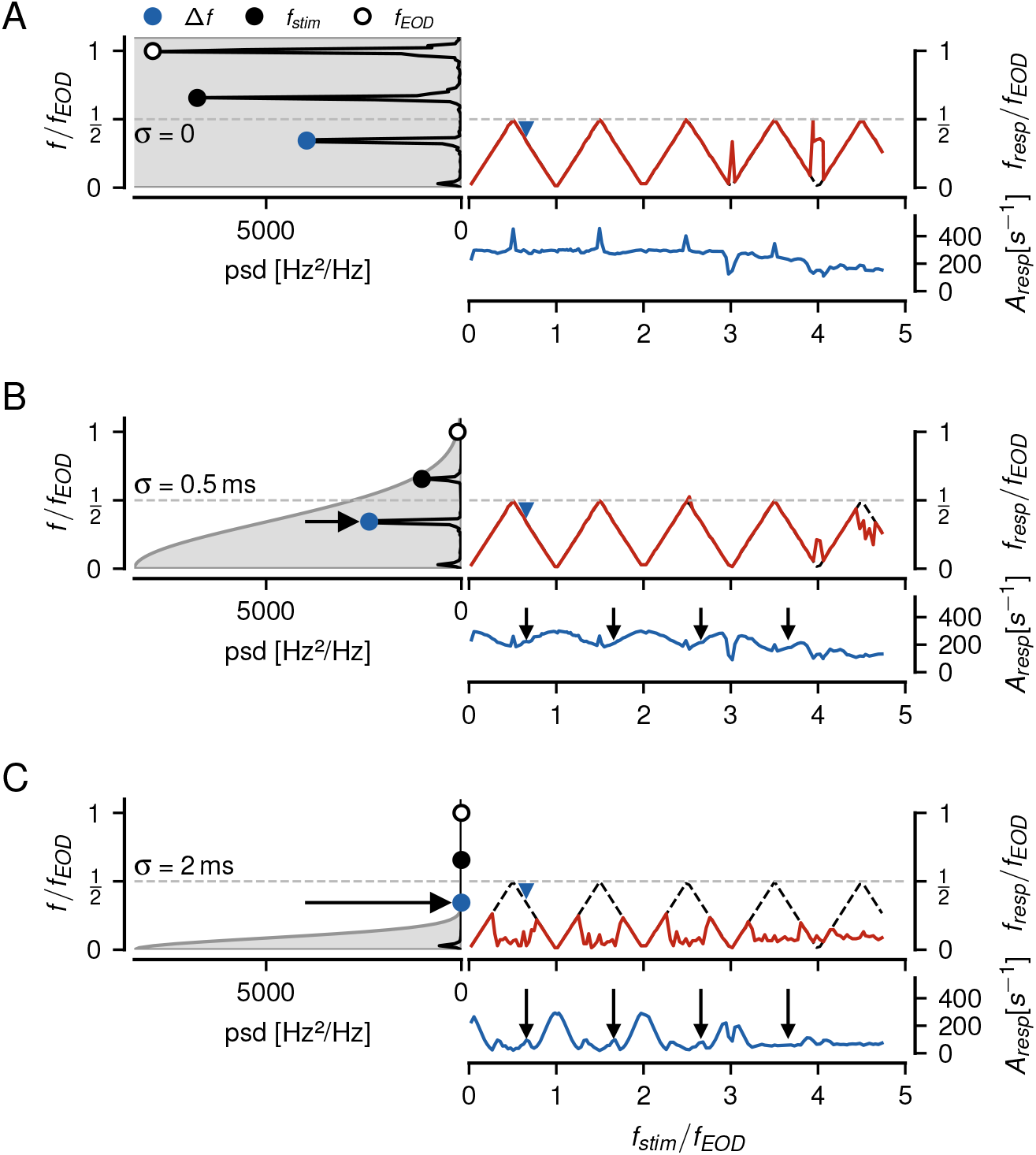
Influence of post-synaptic low-pass filtering on P-unit responses to beat-like stimuli. Left: Power spectra of a P-unit response (same as in Fig. 1) to a Δ*f* = 220 Hz beat (blue markers). The convolution kernel mimicking post-synaptic filtering is indicated by gray area. The horizontal dash-dotted line indicates *f_EOD_*/2. Right: Frequency tuning curve (top) together with the expected aliasing frequencies (black dashed line) and amplitude tuning curve (bottom) estimated from the strongest peak of a P-unit’s response below *f_EOD_*/2. **A** Spectrum and tuning curves of the raw, binary spike trains recorded with a resolution of 40 kHz. The highest peak in the spectrum is at *f_EOD_*(open circle), followed by the peak at the absolute value of the stimulus frequency *f_stim_* (black circle). The peak a the difference frequency corresponding to the frequency of the resulting beat is even smaller, but is the largest peak below *f_EOD_*/2 (orange circle). Frequency tuning follows the aliased frequencies over almost the whole measured range up to 5*f_EOD_*. The amplitude tuning curve is mostly flat with pronounced peaks at odd multiples of *f_EOD_*/2. **B** A biological plausible post-synaptic filter, modeled by convolving the spike trains with a Gaussian kernel (*σ* = 0.5 ms), keeps the frequency tuning, but reduces the amplitude of the P-unit’s response for stimulus frequencies close to odd multiples of *f_EOD_*/2 (arrows). **C** A wider post-synaptic potential, modeled by a Gaussian with *σ* = 2 ms, degrades the frequency tuning curves and strongly modulates amplitude tuning.

### Sensitive cells respond to a larger frequency range

P-unit responses scale with stimulus amplitude (Bastian, 1981; Nelson et al., 1997) and different P-units differ in their sensitivities to a global stimulus (Grewe et al., 2017). To account for both, sensitivity and stimulus intensity, we quantified the P-unit’s response amplitude to a standard stimulus, a beat at Δ*f* = 50 Hz (Grewe et al., 2017). Across all our recorded P-units, firing rate modulations evoked by the 50 Hz beat ranged from 70 to 360 Hz and were positively correlated to baseline firing rates ranging from 80 to 527 Hz (*r* = 0.63, *p*< 0.0001). Think of the modulation depth as the effective stimulus amplitude driving the P-unit’s response.

The stronger the modulation depth of a P-unit’s response, the more the P-unit tuning curves follow the aliasing structure to higher stimulus frequencies (Pearson’s *r* = 0.51, *p* = 0.001, Fig. 3 A). Since we were primarily interested in understanding the mechanisms behind the aliasing structure of the P-units’ responses, we focused our analysis on the *n* = 14 most sensitive cells with modulation depths greater than 265 Hz. These cells respond up to almost four times *f_EOD_* (Fig. 3 B,C) whereas the less sensitive cells responded on average just up to about twice *f_EOD_* (Fig. 3 D,E).

**Figure 3:**
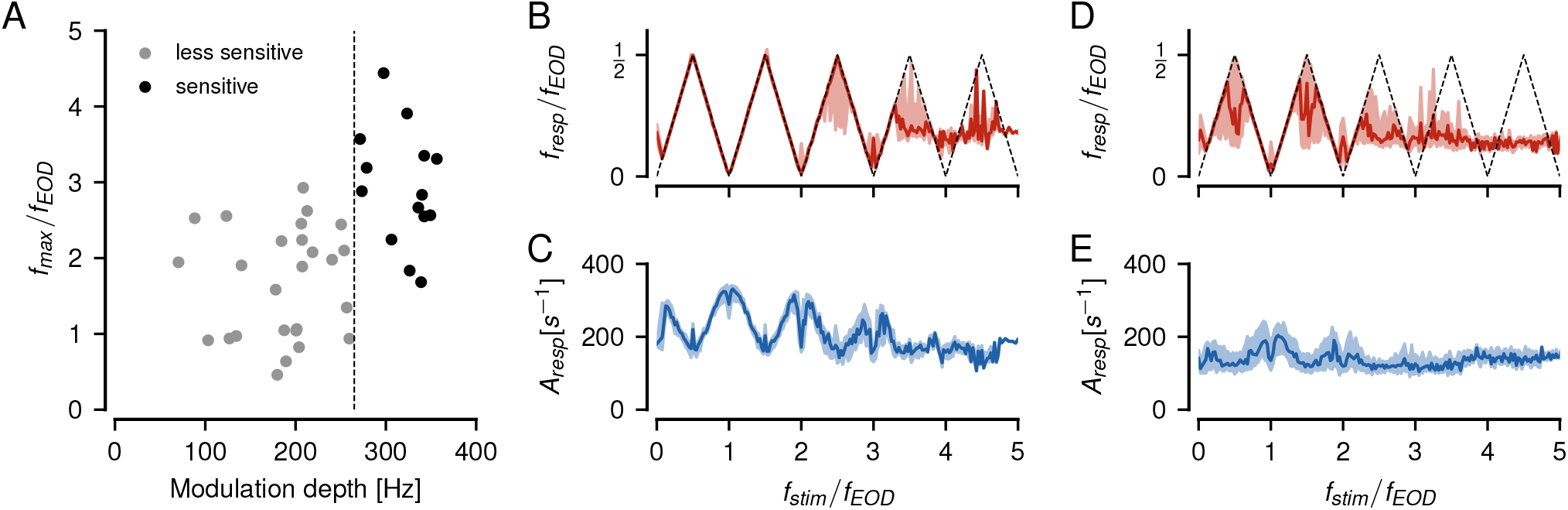
Dependence of the aliasing structure on sensitivity. **A** For each of the recorded P-units we estimated the maximum stimulus frequency *f_max_* up to which its frequency tuning curve indicated a response by following the predicted aliasing curve (e.g. dashed line in panel B). The more sensitive a P-unit, as quantified by the modulation depth of its firing rate response to a 50 Hz standard beat, the higher the maximum frequency. Since we were primarily interested in understanding the mechanisms behind the aliasing structure of the P-units’ responses, we focused our analysis on the most sensitive cells with modulation depths greater than 265 Hz (black). **B** Frequency tuning of the *n* = 14 most sensitive cells reaches up to almost four multiples of *f_EOD_*. **C** Corresponding amplitude tuning curves are of course stronger in comparison to the less sensitive cells. **D, E** Frequency and amplitude tuning curves of the *n* = 26 less sensitive cells (gray dots in A).

### Algorithms for extracting envelope frequencies at high stimulus frequencies

In the following sections we explore prerequisites necessary for neurons to extract the aliased frequency of the amplitude modulations of beats. The resulting theory fully explains the experimental observations, including the enhanced responses at odd multiples of *f_EOD_*/2 (arrows in Figs. 1 J). The details of the mathematical derivations and equations are provided in the supplement.

### Slow beating envelopes in superimposed cosine waves

As suggested by Fig. 1, the aliasing structure of the P-unit response results from the envelopes of the interacting EODs. To understand how such responses can arise, we need to understand the mechanism by which envelopes are retrieved from the superposition of the two EODs. Transformed to the Fourier domain, we ask how a spectral peak at the envelope frequency can be generated. This is a generic problem independent of the electrosensory system, and thus we express the problem in terms of two cosine waves: a carrier, the EOD of the receiving fish, with frequency *ω*_1_ = 2*πf*_1_ and amplitude one and a stimulus, the EOD of another fish, with frequency *ω*_2_ = 2*πf*_2_ and an amplitude *α* measured relative to the amplitude of the carrier (stimulus contrast). As we point out below, we mainly focus on the case *α* ≪ 1, where we have a clear distinction between a large amplitude carrier signal with frequency *ω*_1_ and a stimulus of smaller amplitude at frequency *ω*_2_.

Both signals superimpose:

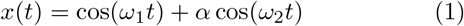

The resulting signal *x*(*t*) also shows the characteristic beating envelopes reoccurring at multiples of the frequency *ω*_1_ of the carrier signal (Fig. 4 A_i_ – A_v_). A running average that attenuates the fast carrier signal, however, results in flat lines, except for stimulus frequencies close to zero (compare Fig. 4 A_ii_ – A_v_ to A_i_). This happens even for *ω*_2_ close to twice and four times *ω*_1_, where upper and lower envelopes are out of phase, because the stimulus cosine distorts the carrier cosine. The respective power spectra have peaks only at the original stimulus frequencies *ω*_1_ and *ω*_2_ (Fig. 4 B_i_ – B_v_). There are no spectral peaks at the slow envelope frequencies (except for *ω*_2_ close to zero), although the envelopes are clearly visible.

**Figure 4:**
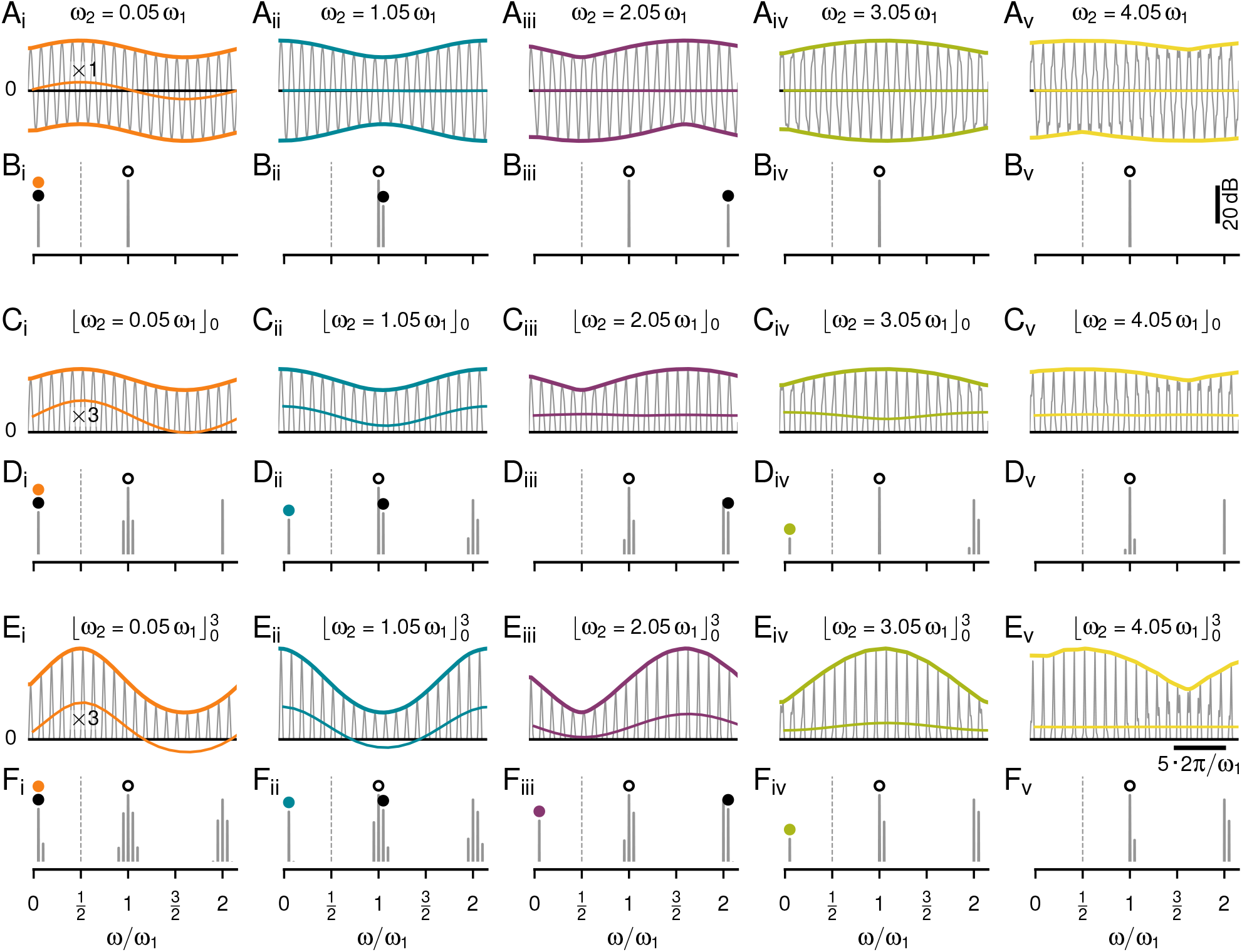
Slow envelopes resulting from the superposition of two cosine waveforms. **A** The raw signals — superpositions of a cosine carrier with fixed frequency *ω*_1_ and stimulus cosines of various frequencies *ω*_2_ close to multiples of *ω*_1_ as indicated and an amplitude of *α* = 0.2 relative to the one of the carrier. Upper and lower envelopes are highlighted by thick colored lines. The thin colored lines are running averages over a single cycle of the carrier frequency, that keep only frequency components smaller than *ω*_1_. For stimulus frequencies similar or larger than the carrier frequency, the running average is zero, even when the upper and lower envelopes are out of phase. **B** Although slow envelopes are clearly visible in the waveforms shown in A, the corresponding power spectra have only peaks at *ω*_1_ (open circle) and *ω*_2_ (black circle) and not at the slow envelopes. Only when the stimulus frequency *ω*_2_ is below *ω*_1_/2 it coincides with the envelope (orange circle in B_i_). **C** Thresholding cuts away the negative half-waves and leaves only the upper envelope intact. Now the running average (enlarged by a factor of three around its mean) follows the envelopes for *ω*_2_ close to 0, 1, and 3 multiples of *ω*_1_, but not for 2 and 4 multiples. Note, for *ω*_2_ = 3.05 *ω*_1_ the running average is antiphasic to the envelope. **D** Thresholding generates frequencies explaining the envelopes at odd multiples of the carrier frequency only (cyan and green circles in B_ii_ and B_iv_). **E** Raising the thresholded signal from C to a power of three narrows the half-waves of the carrier and slightly distorts the envelopes. The running average (enlarged by a factor of three) follows the upper envelope up to three multiples of *ω*_1_, but with decreasing amplitudes. **F** This operation adds a peak in the power spectrum also for stimulus frequencies around the second multiple of the carrier (purple circle in F_iii_).

### Neither the analytic signal nor squaring explains the aliasing structure of the beating envelopes

A non-linear operation needs to be applied to the signal to generate additional spectral peaks at the observed envelope frequencies. Commonly used non-linearities to retrieve the amplitude modulation of a beat for two spectrally close signals are the absolute value of the analytic signal obtained by means of a Hilbert transformation, squaring, or thresholding (Middleton et al., 2007; Stamper et al., 2012).

Both the analytic signal and squaring predict the frequency of the beating envelope to be identical to the difference frequency. This is exactly what we expect for low difference frequencies, i.e. for stimulus frequencies *ω*_2_ close to *ω*_1_. For higher difference frequencies, however, the analytic signal suggests an amplitude modulation with growing frequency (Fig. S1 A), although the signals are not necessarily amplitude modulated signals anymore (e.g. panels A_i_, A_iii_ and A_v_ in Fig. 4, upper and lower envelopes are no mirror versions of each other). Squaring also predicts a spectral peak at the difference frequency, no matter how large the difference frequency (Fig. S1 B). These two types of non-linearities neither explain the aliasing structure we observe in superimposed cosines (Fig. 4 A), nor in superimposed EODs and the corresponding P-unit responses (Fig. 1).

### Thresholding explains aliasing at odd multiples of the carrier frequency

A threshold non-linearity

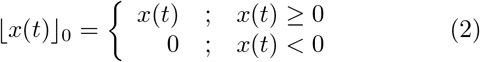

sets all negative values of a signal to zero. Only the positive half-waves are passed through. The upper envelope is retained whereas the lower envelope of the signal is discarded (Fig. 4 C). For brevity we call this half-wave rectification “thresholding”. Thresholding indeed generates spectral peaks in the resulting signal at some of the envelope frequencies, but not for stimulus frequencies close to the second or fourth multiple of the carrier frequency (Fig. 4 D). Consequently, low-pass filtering the thresholded signals results in flat lines for *ω*_2_ close to two and four times of *ω*_1_, despite the obvious envelope. Surprisingly, the stimulus distorts the carrier such that for *ω*_2_ close to 2*ω*_1_ the running average generates a signal with the same frequency as the envelope, but antiphasic to the envelope (Fig. 4 C_iv_).

To make the threshold operation, Eq. (2), analytically tractable, we approximate it by a multiplication of the signal (Fig. 5 A) with a pulse train of the same frequency *ω*_1_ as the carrier signal (Fig. 5 B):

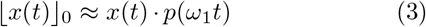

The pulse train

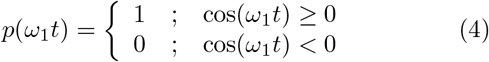

multiplies positive half-waves of the carrier cosine with one and negative half-waves with zero (Fig. 5 C).

**Figure 5:**
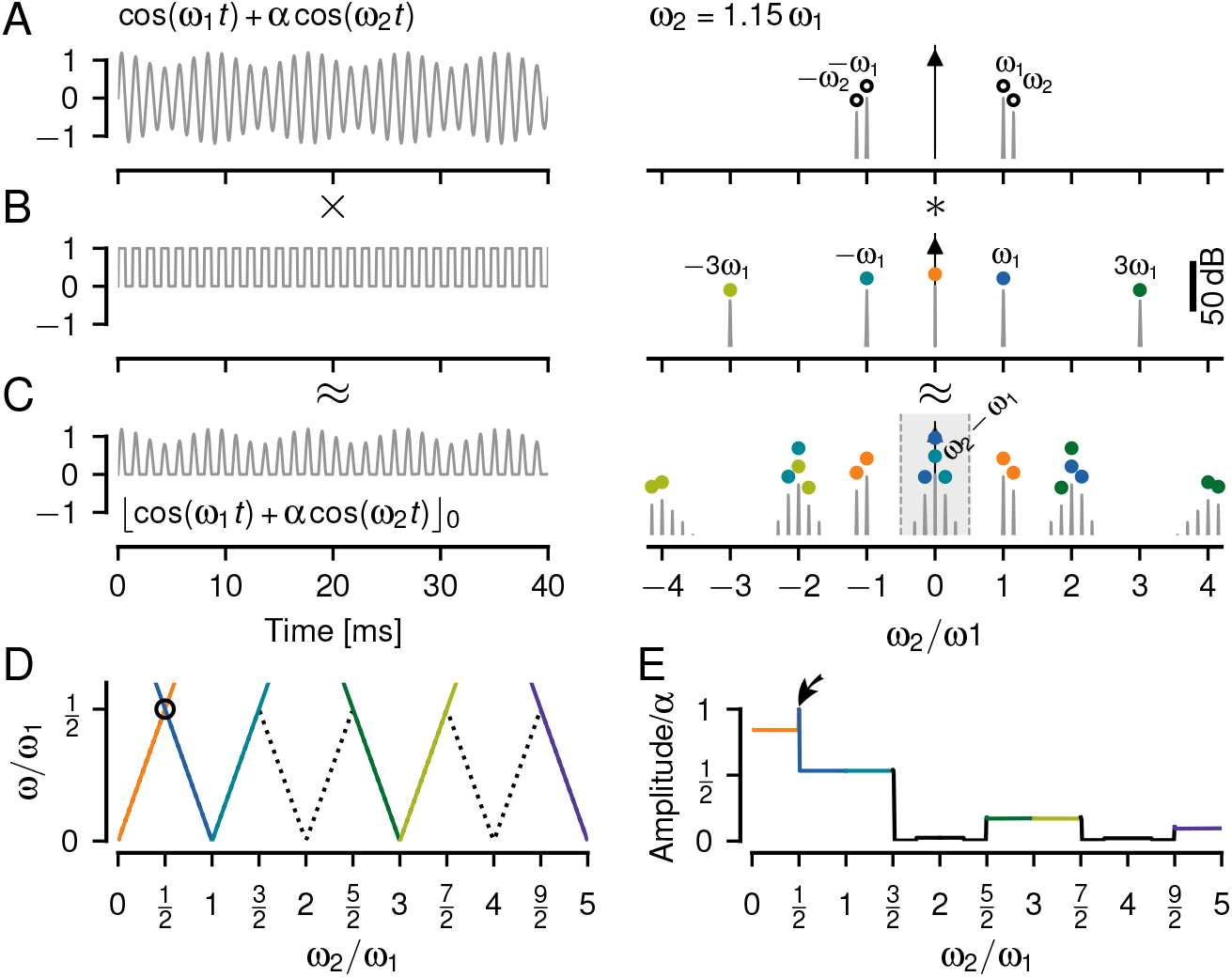
Thresholding is not sufficient to explain the full aliasing structure of beat-like envelopes. **A** A beat generated by summing up two cosines with frequencies *ω*_1_ and *ω*_2_ (left) only contains these two frequencies in its Fourier spectrum (right). The amplitude *α* = 20 % of the stimulus cosine is smaller than the one of the carrier. **B** A pulse train, Eq. (4), used to approximate a threshold operation, Eq. (2), has peaks at odd multiples of *ω*_1_ and at zero. **C** Thresholding the signal from A results in a rich spectrum. In the limit of vanishing stimulus amplitude *α* the thresholded signal can be approximated by multiplying the signal from A with the pulse train in B. The corresponding spectrum is approximated by convolving the spectrum of the signal with the one of the pulse train (colored circles, one color for each peak of the pulse train). Here, for *ω*_2_ = 1.15 *ω*_1_ a peak appears at the difference frequency *ω*_2_ − *ω*_1_ below *ω*_1_/2 (gray area). This peak describes the slow amplitude modulation of the beat visible in A. Because the stimulus amplitude is not close to zero, this approximation does not explain all the side peaks in the spectrum. **D** The position of peaks in the spectra of thresholded signals (colored lines) below *ω*_1_/2, as a function of stimulus frequency *ω*_2_. This curve is a prediction for the frequency tuning curves of P-units. The dashed line marks the expected aliased frequencies, and the circle the only crossing of spectral peaks. Note, that close to two and four multiples of *ω*_1_ no peaks are created by thresholding below *ω*_1_/2. See Fig. S1 C for spectral peaks also at higher frequencies. **E** Amplitude of the peaks shown in D decrease with higher multiples of *ω*_1_. At the crossing of spectral peaks (circle in D) amplitudes sum up (arrow). These amplitudes are measured relative to the amplitude *α* of the stimulus cosine and would drive the amplitude tuning curves of P-units.

Note that this approximation is valid only in the limit *α* → 0. For larger stimulus amplitudes the stimulus distorts the carrier, shifting also its zero crossings. As a consequence, additional side-peaks occur in the spectra (Fig. 4 D).

According to the convolution theorem a multiplication in time equals a convolution in the Fourier domain. Thus, the Fourier spectrum of the thresholded signal (Fig. 5 C) is given by the convolution of the spectrum of the signal (Fig. 5 A) and that of the pulse train (Fig. 5 B). The spectrum of the pulse train has a peak at zero frequency and peaks at all odd multiples of the carrier frequency *ω*_1_.

A first component of the resulting spectrum is the convolution of the carrier frequency, *ω*_1_, with the pulse train spectrum. This results in peaks at even multiples of *ω*_1_ and at ± *ω*_1_ (horizontal lines in Fig. S1 C). These frequency components do not make up the beating envelope, because they do not depend on *ω*_2_.

The second component of the spectrum, the convolution of the stimulus frequency *ω*_2_ with the pulse train, provides side peaks at ±*ω*_2_ to all the peaks of the pulse train (Fig. S1 C). These peaks explain the aliasing structure of the beating signal envelopes and thus the frequency tuning curves of P-units’ around odd multiples of *ω*_1_ and around zero frequency, but not around two times *ω*_1_ (Fig. 5 D). The amplitudes of these peaks quantify the amplitudes of the beating envelopes and drive the amplitude tuning curves of P-units. They decrease with higher multiples of the carrier (Fig. 5 E, see supplement for a mathematical derivation of these amplitudes). Note that this decrease in amplitude is stronger than that of the signal’s envelopes.

In contrast to the amplitude of the analytic signal and to squaring, the threshold operation introduces many additional peaks in the spectrum. These are necessary for explaining some but not all of the aliasing structure of signal envelopes. In particular, thresholding does not generate low frequency spectral peaks for stimulus frequencies around twice the carrier frequency.

### Threshold cubed fills in frequencies at around twice the carrier frequency

How can we fill in the missing components in the spectrum around even multiples of *ω*_1_? Let’s try to make the threshold operation more non-linear by raising its output to a power of three:

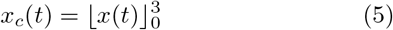

In the resulting signals the half-waves of the carrier are more narrow and the envelopes are slightly distorted in comparison to the pure threshold (Fig. 4 E). In the corresponding power spectra we now get peaks at the slow envelope frequencies up to the third multiple of *f_EOD_* (Fig. 4 F). In particular, we get such a peak for stimulus frequencies close to the second multiple of *f_EOD_* (Fig. 4 F_iii_). Also the running average produces signals of the same frequency and phase as the envelopes up to the third multiple of *f_EOD_*.

Again, this can be approximated by multiplying with a pulse train after taking the signal to the power of three (Fig. 6 A–C). The spectrum of the two superimposed cosines cubed has 2^3^ = 8 peaks (two times convolution of two peaks with themselves, Fig. S2 A). Of those the peaks at |2*ω*_1_ − *ω*_2_| = |*ω*_1_ − Δ*ω*| are the only relevant additions in comparison to the threshold without exponent. The convolution of these peaks with the zero-frequency peak of the pulse train fills in the missing frequencies around twice the carrier frequency (Fig. 6 D, red and purple). This is enough for explaining the frequency tuning curve of P-units (Fig. 3 B) up to 3.5 multiples of *f_EOD_*.

**Figure 6:**
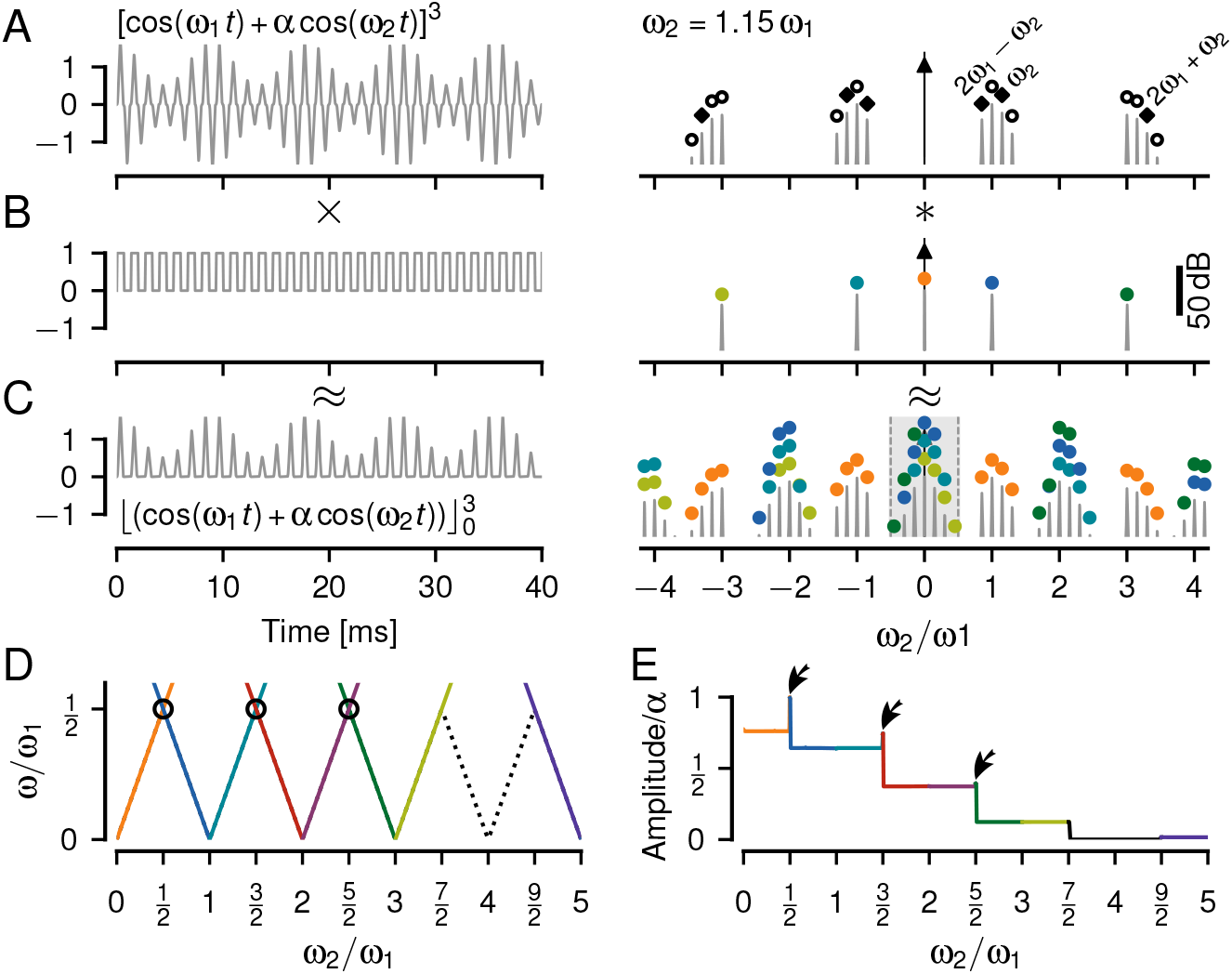
Cubing fills in frequencies corresponding to slow envelopes close to the second multiple of the carrier. **A** The cubed signal (left) has a more pointed carrier waveform and eight peaks in the positive half of the Fourier spectrum (right, Fig. S2 A). **B** Again we approximate the threshold operation by a multiplication with a pulse train. **C** The threshold-cubed signal has a much richer power spectrum compared to the thresholded signal without exponent (Fig. 5 C). The convolution process is illustrated by the colored markers — the spectrum of the cubed signal from A is shifted to each of the individually colored peaks in the spectrum of the pulse train from B. In this process many peaks fall on top of each other and add up. The gray shading marks frequencies below *ω*_1_/2. **D** Position of the largest spectral peaks of thresholded and cubed signals as a function of stimulus frequency. This is the frequency tuning curve. See Fig. S2 B for spectral peaks at higher frequencies. Dashed line indicates the expected fully aliased frequencies, that are extracted by the cubed-threshold operation up to *ω*_2_ = 3.5 *ω*_1_. Circles mark peak crossings. **E** Amplitudes of the peaks in D describe the amplitude of the beat-like signal envelopes. Crossing spectral peaks in D (circles) add up and result in elevated amplitudes (arrows). These amplitudes are driving the P-unit responses and thus are relevant for their amplitude tuning curve.

The amplitudes of the peaks below *ω*_1_/2 decline in a step-wise manner for each multiple of *ω*_1_ (Fig. 6 E). At *ω*_1_/2, 3*ω*_1_/2, and 5*ω*_1_/2 spectral peaks cross each other (circles in Fig. 6 D). Their respective amplitudes add up and result in elevated amplitudes exactly at these frequencies (Fig. 6 E, arrows). These peak crossings explain the elevated P-unit responses at these frequencies (arrows in Fig. 1 J). See supplement for a mathematical derivation of peak frequencies and amplitudes and Fig. S2 B. Again, the peak amplitudes decline much faster than the amplitudes of the envelopes.

In conclusion, a cubed threshold operation is sufficient to generate spectral peaks below half the carrier frequency for stimulus frequencies up to inclusively three multiples of *ω*_1_. These peaks correspond to the aliasing structure we observe in the envelopes of these signals and potentially explain the recorded P-unit responses. In contrast, a pure threshold fails to extract slow envelopes around twice the carrier frequency. In addition, the non-linear operation needs to be followed by a low-pass filter that isolates these envelope frequencies by attenuating the carrier frequency and other spectral peaks beyond *ω*_1_/2.

### Spiking dynamics cannot explain responses to higher beat frequencies

Eventually, a spike generator, here modeled as a leaky integrate-and-fire neuron (LIF) with adaptation current, encodes the extracted envelope in a train of action potentials (Fig. 7). LIF models with a threshold non-linearity without exponent (power of one), Eqs. (A.1), (A.3), (A.4) (Sinz et al., 2020), were individually fitted to baseline and step-response characteristics of *n* = 9 recorded cells.

**Figure 7:**
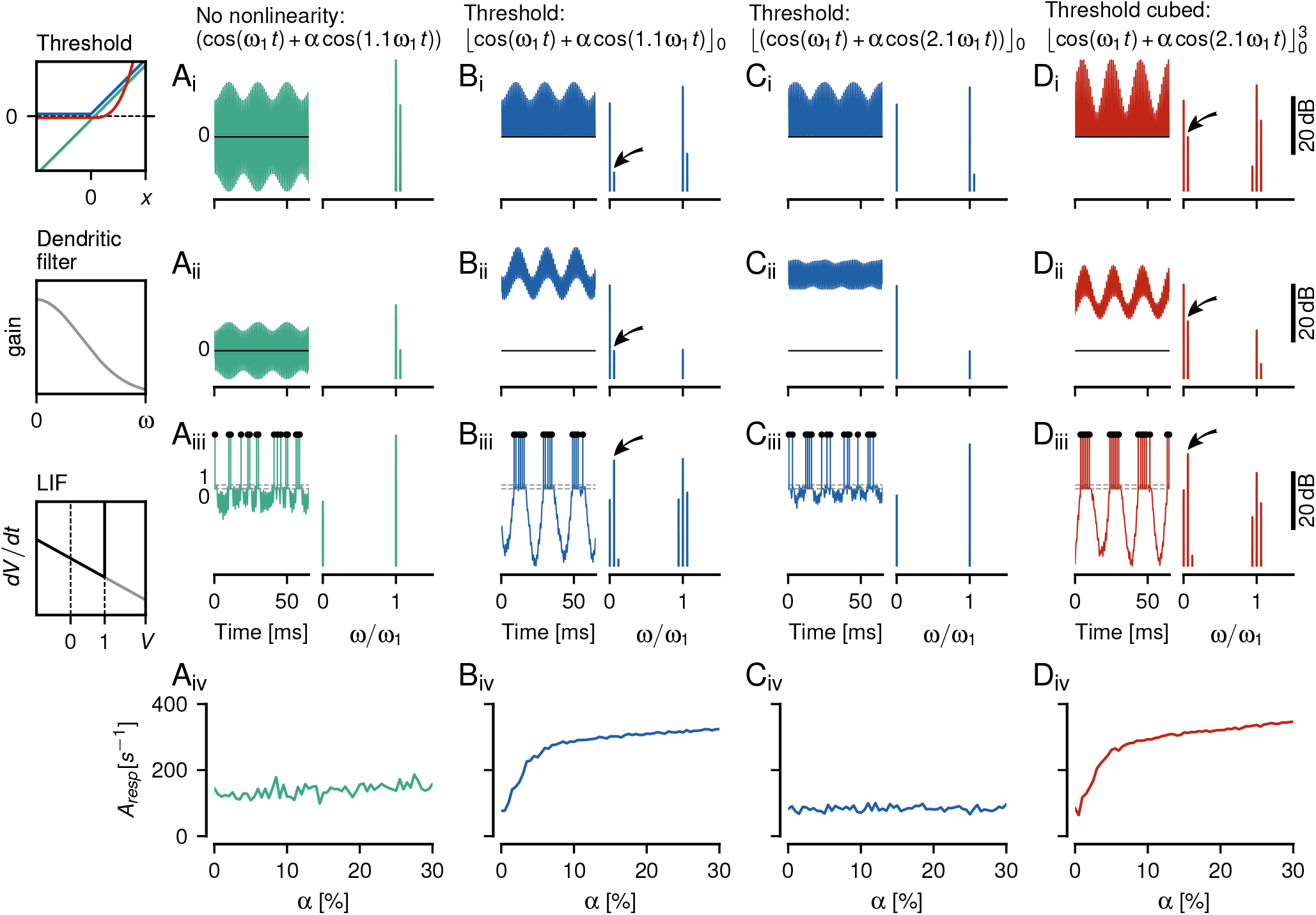
Integrate-and-fire models of P-unit spiking activity. As an example, simulations of the model for cell 2018-06-25-ad are shown (see Tab. A.1 for parameters). **A**_i_ Waveform (left) and corresponding power spectrum (right, same reference for decibel scale for all spectra shown in top two rows and a different reference for the spectra of the third row) of two superimposed cosine waves. No non-linearity is applied (green identity line in box). **A**_ii_ Passing the signal through a dendritic low-pass filter, Eq. (A.4), here *τ_d_* = 1.88 ms, attenuates the two frequencies. **A**_iii_ A leaky integrate-and-fire neuron (LIF) does not encode the beating amplitude modulation in its spike train. **A**_iv_ The height of the spike-train spectrum at the beat frequency grows weakly with beat amplitude *α*. **B** Applying a threshold (blue curve in threshold box) generates a peak in the power spectrum at the beat frequency (arrow), if the stimulus frequency *ω*_2_ is close to the carrier frequency *ω*_1_. This frequency component becomes more apparent after dendritic low-pass filtering and the LIF is able to generate spikes at the peaks of the beat. The spike times contain a spectral peak at the beat frequency (arrow) and the size of this peak strongly depends on beat amplitude. **C** For a stimulus frequency close to twice the carrier frequency, however, a threshold does not produce a spectral peak that would represent the beating envelope. After the dendritic low-pass filter the waveform still oscillates symmetrically around a fixed mean value and the LIF responds with tonic spiking that is not modulated by the signal’s envelope. **D** A cubed threshold (red curve in threshold box), however, generates a peak at the envelope frequency (arrows). Dendritic low-pass filtering results in an oscillating signal that shifts up and down according to the signal’s envelope. Consequently, the LIF is able to encode this envelope in its spiking activity and is also sensitive to the amplitude of the envelope.

The non-linear spiking dynamics on its own is not sufficient to extract and respond to a beating amplitude modulation resulting from two spectrally close frequencies. Although some models generate a peak at the beat frequency, their responses are almost independent of stimulus amplitude (Fig. 7 A).

A threshold non-linearity generates a sufficiently large spectral peak at the difference frequency. The spike generator is driven by this frequency and generates action potentials that encode the beat. Thresholding the signal results in modulations of the evoked firing rate responses that depend strongly on beat amplitude, faithfully reproducing the spiking properties of P-units (Fig. 7 B).

For stimulus frequencies close to twice the carrier frequency, however, action potentials do not encode the apparent envelope of the signal, because after thresholding and low-pass filtering, the input to the spike generator does not provide a frequency component at the envelope frequency as input (Fig. 7 C).

As worked out above, it requires a power of three applied to the thresholded signal to make the spike generator respond to the envelope at stimulus frequencies close to twice the carrier frequency (Fig. 7 D). Neither the hard threshold non-linearity the LIF applies on the membrane voltage nor the smooth voltage-threshold, of the exponential integrate-and-fire neuron (EIF, data not shown, Fourcaud-Trocmé et al., 2003) can replace the power of three to extract the envelope resulting from a stimulus at around twice the carrier frequency. The non-linear dynamics of a spike generator can not generate the aliasing structure of the P-unit responses. Rather, a sufficiently strong static non-linearity (Fig. 7 D_i_) has to be applied to the signal, such that the necessary low-frequency peak in the spectrum is generated. Subsequent low-pass filtering then isolates this peak (Fig. 7 D_ii_) and this is what the spike generator responds to.

### A power of three describes P-unit responses best

For a more systematic evaluation which exponent on the threshold operation describes the P-unit responses best, we simulated the LIF models using exponents at the threshold non-linearity ranging from *p* = 0.2 to 5. The resulting frequency and amplitude tuning curves were compared to the experimentally measured ones (Fig. 8).

**Figure 8:**
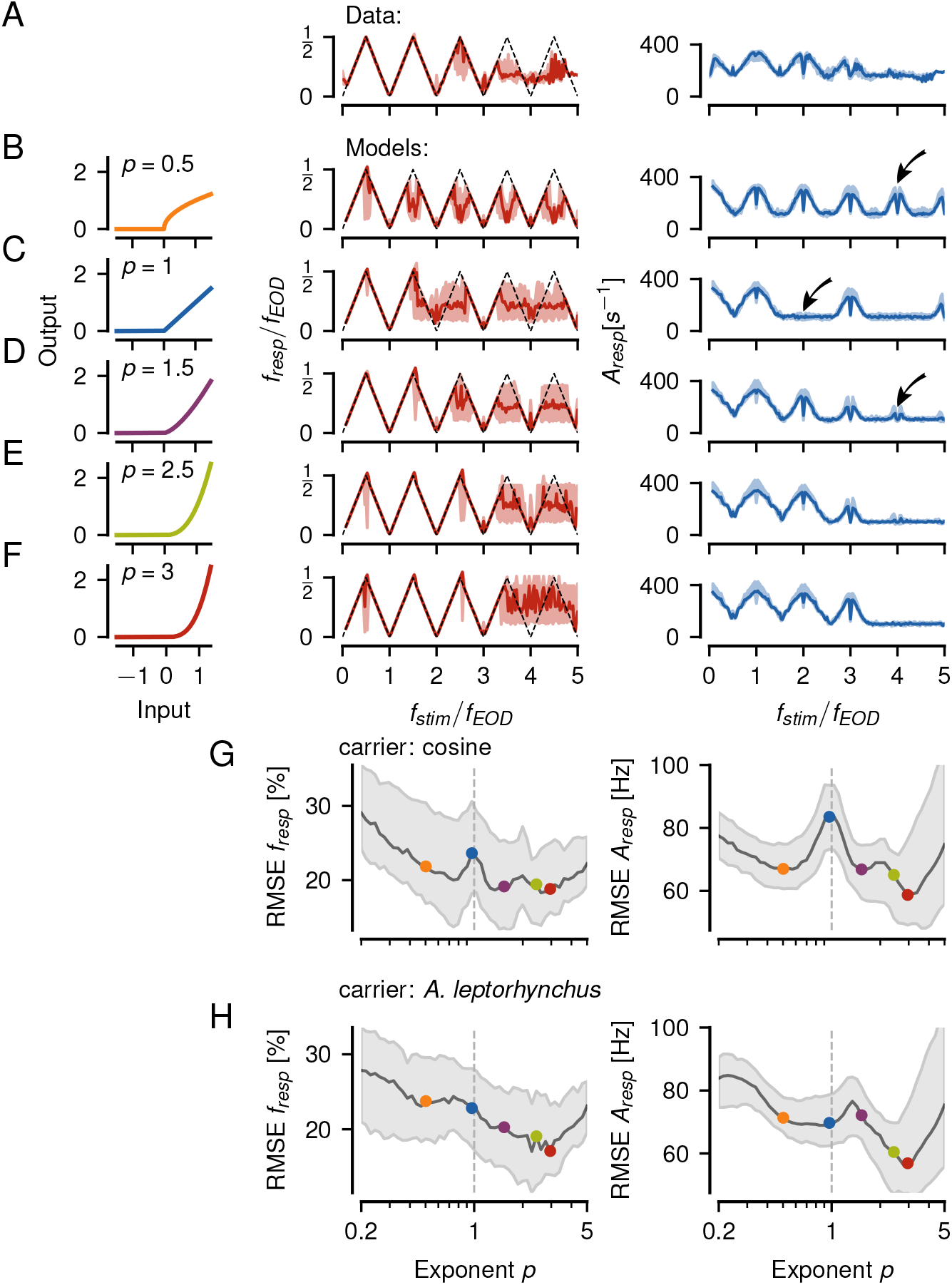
Dependence of LIF model performance on threshold exponent. **A** Frequency (left) and amplitude tuning curves (right, population medians with interquartile ranges) of *n* = 14 experimentally measured P-units of highest sensitivity (same as Fig. 3 B,C). **B–F** Tuning curves simulated from a population of *n* = 9 LIF models (Fig. 7) that have been fitted to individual P-units. Exponent *p* of the threshold non-linearity applied to the signal as indicated and illustrated in the left column. Arrows indicate missing or additional responses in the amplitude tuning curve. **G** Root mean squared error (RMSE, median with interquartile range) between the tuning curves of each experimentally measured P-unit shown in A and each LIF model in dependence on the threshold exponent used in the models. Examples from B–F are indicated by the correspondingly colored circles. Here, both the carrier and the stimulus are pure sine waves. **H** Same as in G but with an EOD waveform of *A. leptorhynchus* as carrier and a pure sine wave as stimulus, resembling the situation in the electrophysiological experiments (see Fig. 9).

As predicted, a pure threshold without exponent shows responses at the zeroth, first and third harmonic, but diverges from the measured activity (Fig. 8 A) around the second harmonic (Fig. 8 C). Exponents both higher (Fig. 8 D, E, F) and lower than one (Fig. 8 B) fill in the response at the second multiple of *f_EOD_*. Models with powers of 0.5 and 1.5 additionally respond to the forth multiple of *f_EOD_*.

To quantify the model performance we computed root mean squared errors (RMSE) between all pairings of the 14 experimentally measured cells with the 9 model cells of similar sensitivity. The RMSEs between frequency tuning curves were minimal at powers of about 0.8, 1.5, and 3 (Fig. 8 G, left). The RMSEs for the corresponding amplitude tuning curves showed similar minima but with the smallest RMSE at a power of three (Fig. 8 G, right). A power of three indeed describes both the frequency and amplitude tuning curves of P-unit responses best.

### Harmonics of the carrier are not sufficient to explain aliasing

So far our reasoning was based on pure sine waves. In the electrophysiological recordings, however, the carrier was a real EOD waveform of *A. leptorhynchus*. Using these EOD waveforms instead of sine waves for the models made the minimum at an exponent of *p* = 3 more distinct (Fig. 8 H). In a model with a pure threshold (*p* = 1) the harmonics of the carrier do not contribute to shift the stimulus frequency to all multiples of *f_EOD_*. A wider or narrower EOD waveform, however, modifies the aliasing structure introduced by the threshold operation in a way we do not observe in the data. Adding a power of three to the threshold makes the P-unit responses more robust against changes in the EOD waveform (Fig. 9, see sections 2.6 and 2.7 in the supplement for a more detailed explanation).

**Figure 9:**
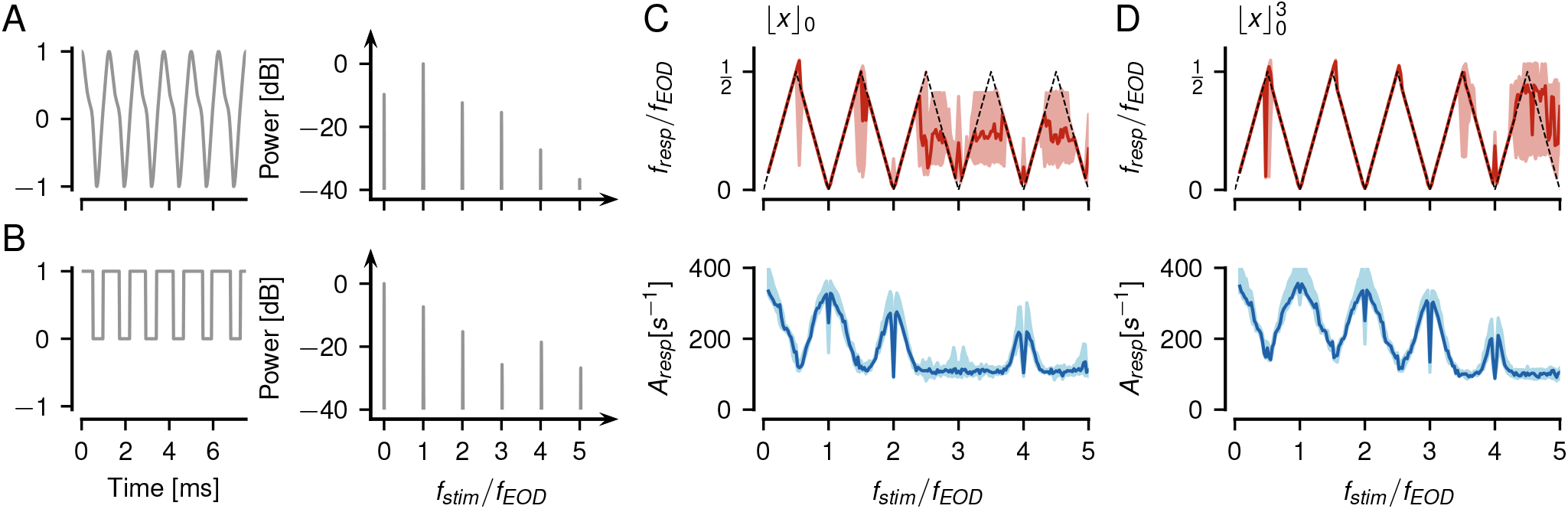
Influence of higher harmonics of the carrier on aliasing. **A** An EOD waveform and the corresponding power spectrum of an *A. leptorhynchus* used as a carrier signal of frequency *ω*_1_ for simulating P-unit responses with the LIF models from Fig. 8. **B** The corresponding pulse train (left), needed to approximate the threshold operation, Eq. (2), has a duty cycle larger than 50 %. Its spectrum (right) has a peak at the second multiple of *f_EOD_* in contrast to the spectrum of a pulse train with a 50 % duty cycle (Fig. 5 B). **C** The enlarged duty cycle modifies the aliasing pattern of the resulting frequency (top) and amplitude (bottom) tuning curves, when using a threshold without exponent. For the example EOD waveform shown, the third harmonics and not the second as for a pure cosine wave is missing. **D** Taking in addition the thresholded signal to a power of three makes the tuning curves more independent of the duty cycle, an in particular fills in responses around three multiples of *f_EOD_*.

### Ambiguous beats evoke similar behavioral responses

Our results imply that P-unit responses are potentially ambiguous with respect to the absolute stimulus frequency, because P-unit responses to similarly mistuned multiples of *f_EOD_* differ only in modulation depth of their firing rate responses. We tested this hypothesis behaviorally by means of the jamming avoidance response (JAR, Watanabe and Takeda, 1963). When a receiving fish is stimulated with a sinusoidal mimic that is close to but below the own EOD frequency, it will raise its EOD frequency by a few Hertz on a timescale of about 10 seconds (Fig. 10 A). We repeated the experiment with stimulus frequencies 5 Hz below one to five times *f_EOD_*. Indeed, all the fish tested (*n* = 5) responded with a significant increase of their EOD frequency to stimulus frequencies close to one, two, and three multiples of their EOD frequency (Fig. 10 B). None of the fish responded to four times *f_EOD_*, but some fish slightly elevated their EOD frequency in response to five times *f_EOD_* by less than 1 Hz. The size of the frequency shift in response to non-zero multiples of *f_EOD_* approximately follow the amplitudes of the corresponding envelopes predicted by a cubed threshold (Fig. 10 C). To the stimulus at the zeroth multiple of *f_EOD_* only a single fish responded with a noticeable frequency shift.

**Figure 10:**
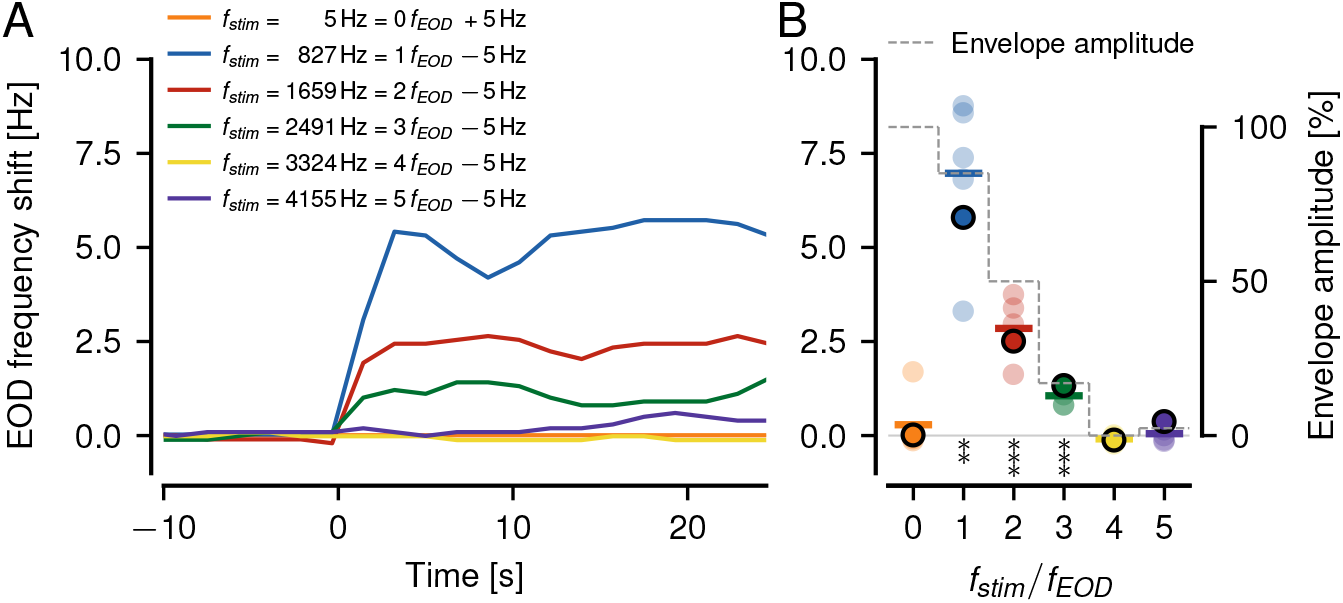
Jamming Avoidance Response (JAR) to stimuli close to multiples of *f_EOD_*. **A** Shifts in EOD frequency of a single example fish induced by sinusoidal stimuli switched on at 0 s. The stimulus frequency was set 5 Hz above or below 0 to 5 times the EOD frequency of the fish as indicated. The fish responded to this stimulus by shifting its EOD frequency within about 10 s to a higher value. **B** Steady-state frequency shifts measured between 15 and 25 s after stimulus onset of five fish as a function of *f_EOD_* multiple. Frequency shifts of the fish shown in A are highlighted by circles with black outline. The frequency shifts averaged over the five fish (colored horizontal lines) approximately follow the corresponding envelope amplitudes predicted by a cubed threshold (dashed line, see supplement section 2.4 for equations) for non-zero multiples of *f_EOD_*. Frequency shifts in response to low absolute stimulus frequencies are mostly absent, despite having the largest envelope amplitude. Asterisks indicate significant deviations of the steady-state frequency shifts from zero (t-Test, * * *p*< 1 %, * * \**p*< 0.1 %)

## 3. Discussion

We observed that P-units encode a wide range of stimulus frequencies up to about 3000 Hz (Fig. 1). Superposition of the EOD carrier and the stimulus led to beating signal envelopes. At integer multiple of the carrier frequency these envelopes are slow and between integer multiples they are fast, a pattern reminiscent of aliasing known from the sampling theorem. In accordance with reoccurring envelopes, the P-unit tuning curves are also periodic in multiples of the EOD frequency, while the amplitude of the responses declined with higher stimulus frequencies.

### P-unit tuning to high difference frequencies

P-unit tuning curves for beat stimuli have so far been only measured for difference frequencies up to about 300 Hz (Bastian, 1981; Nelson et al., 1997; Benda et al., 2006; Walz et al., 2014). The amplitude of the firing-rate modulation induced by beating envelopes resembles a band-pass tuning: At frequencies close to zero the P-unit responses are reduced (Nelson et al., 1997) which reflects the high-pass filter induced by fast spike-frequency adaptation (Benda et al., 2005). Towards higher beat frequencies the response was found to steadily decline towards zero as expected for a spiking neuron (Fourcaud-TrocmÉ et al., 2003) and as set by the neuron’s baseline firing rate (Knight, 1972).

The experimental findings reported here clearly demonstrate that P-units do show responses to difference frequencies beyond 300 Hz. Instead of the expected steady decline, the tuning repeats at multiples of the EOD frequency. This has profound consequences for the encoding of electrocommunication signals, so-called chirps, that transiently change the difference frequency (Benda et al., 2005, 2006). The P-unit response to chirps could be explained by transient firing rate modulations mediated by the P-units’ amplitude tuning curve (Walz et al., 2014). In a field study the behavioral relevance of chirps at difference frequencies beyond 400 Hz and thus beyond *f_EOD_*/2 of the female has been observed (Henninger et al., 2018). A monotonously declining tuning curve would not suffice to explain how chirps could be encoded at such high difference frequencies. The repetitive tuning we describe here retains the capability to change the modulation depth of the firing rate in response to chirps.

In this context, the low-pass filtering happening at the synapse between afferents and their targets, the pyramidal cells in the electrosensory lateral line lobe, plays an important role (Fig. 2). If the kernel is too narrow, then the tuning curve of the P-units is almost flat and firing rate modulations caused by chirps would be quite small.

If the kernel is too wide, the tuning curve is only modulated within a narrow range of stimulus frequencies around multiples of the carrier frequency. Only for kernels resembling the experimentally measured postsynaptic potentials (*σ* = 0.5 ms, Berman and Maler, 1998) is the tuning curve fully modulated without being flat between the multiples (Fig. 3). We therefore hypothesize electric fish with lower EOD frequencies, like for example *Eigenmannia* spec., to have correspondingly wider postsynaptic potentials.

Surprisingly we observed P-unit responses down to absolute stimulus frequencies of about 35 Hz, a range commonly assumed to be primarily driving the ampullary electrosensory system (Kalmijn, 1974; Engelmann et al., 2010; Grewe et al., 2017). Towards even lower stimulus frequencies responses vanish, although there seems to be a large variability between cells (not shown). Future recordings with finer resolution of the stimulus frequencies are needed for in depth exploration of this specific frequency range.

### Envelope extraction at high difference frequencies

The repetitive tuning curve of P-units is in a way trivial in that it simply follows the envelopes visible in the superimposed cosine signals. But how are these envelopes extracted from the original signals, that do not contain the envelope frequencies in their power spectra?

Usually, the envelope frequency is considered equal to the difference frequency (Walz et al., 2014; Joris et al., 2004). We have shown that extracting the envelope by the Hilbert transform or squaring of the signal (Middleton et al., 2007; Longtin et al., 2008; Stamper et al., 2012) fails for higher difference frequencies and are not sufficient to explain the experimental observations (Fig. S1 A, B). Both lead to spectral peaks that reflect the true envelope for frequency differences below half the carrier frequency but fail beyond. Furthermore, for stimulus frequencies close to even multiples of the carrier frequency, the resulting signals cannot be described as amplitude modulations, because their lower and upper envelope are out of phase (Fig. 4 A).

For analyzing EEG/EMG (Myers et al., 2003) or acoustic signals (Khanna and Teich, 1989) simple thresholding is often applied to compute envelopes. Such a threshold, in deep learning also known as a Rectifying Linear Unit (ReLU), extracts envelope frequencies only at odd harmonics of the carrier (Fig. 5). Our results show that a threshold operation followed by exponentiation is required to explain the experimental findings (Figs. 6 and 8). Neither the non-linearity of action-potential generation (Fig. 7), nor higher harmonics of the carrier and the signal are sufficient to substitute the exponentiated threshold (Fig. 9).

Interestingly, it has become common to apply smooth threshold functions such as ELU (Clevert et al., 2015) or Softplus (Glorot et al., 2011) in deep learning approaches using artificial neural networks. All of these are potential alternatives for the threshold raised to a power of three we are suggesting here, and can be approximated by a ReLU raised to a power of three in the vicinity of their threshold. The same holds true for sigmoidal activation functions which are discussed for the transformation of a hair-cells membrane voltage by their ribbon synapse (Peterson and Heil, 2019). Only for larger inputs their saturation will lead to noticeable deviations.

### Extraction of secondary envelopes

Secondary envelopes, the modulation of the amplitude of beats, arise from relative movements between two fish (Yu et al., 2005) as well as interactions between more than two fish (Middleton et al., 2006; Stamper et al., 2012). They provide a context onto which electrocommunication signals are encoded in the thalamus (Wallach et al., 2022). Secondary envelopes, however, are mainly extracted downstream of P-units in the ELL (Middleton et al., 2006) by means of threshold non-linear response curves of the involved neuron (Middleton et al., 2007). The input from which the envelopes are extracted are not superpositions of sine waves anymore, but rather temporally modulated population firing rates. Therefore, the problem of encoding high stimulus frequencies does not exist in the context of encoding secondary envelopes. However, our results demonstrate that secondary envelopes are also to be expected on slow beating envelopes arising close to multiples of the carrier frequency and that these will also be encoded in the electrosensory system.

### Sinusoidal amplitude modulations (SAMs) versus beats

SAMs of various frequencies have been used to characterize signal processing in the electrosensory (Bastian, 1981) as well as the mammalian auditory system (Joris et al., 2004). SAM stimuli multiply a carrier – the EOD of an electric fish or a tone — with a periodic amplitude modulation, like in Eq. (S6), and differ from super-imposed cosines, Eq. (1), by having three spectral peaks instead of two. The additional side-peak of a SAM stimulus already fills in responses at the second multiple of the carrier frequency when used in conjunction with a threshold without exponent (Fig. S2 C). This effect would have obscured the necessary cubed threshold, if we had used SAMs instead of realistic superimposed cosines.

### Relation to the sampling theorem

In the limit to an infinitely high exponent, the thresholded and exponentiated carrier approaches Dirac delta functions positioned at multiples of the carrier’s period. This pulse train can be thought to sample the stimulus waveform with the carrier frequency exactly like in the setting of the sampling theorem. However, the stimulus waveform also gets transformed by the threshold and the exponentiation. The higher the exponent the larger the distortion of the extracted envelope. Thus, the exponent should not be too large in order to maintain an accurate representation of the amplitude modulation. In this sense, a sharp threshold without exponent would be ideal.

### Physiological mechanisms for beat extraction

Our theoretical considerations suggest an exponent of at least three. What are possible physiological substrates of such a non-linear operation?

First, we need to note that a perfect threshold is a mathematical abstraction. Any physiological mechanism implementing a threshold operation has a smooth transition. A cubic power applied to the threshold approximates such a smooth threshold (Fig. 7 D) and in that sense can be considered biologically more plausible than a perfect threshold.

The most likely site for the smooth threshold operation are the ribbon synapses of the electroreceptor cells onto their afferents (Szabo, 1965; Wachtel and Szamier, 1966) as has been suggested previously (Chacron et al., 2001). The sigmoidal shape of the activation function of voltage-gate calcium channels that trigger synaptical transmitter release, for example, implement a threshold operation. Unfortunately, the synapse connecting primary electroreceptors and the afferents are difficult to access and there are no recordings of their transfer function.

One afferent receives input from a set of primary electroreceptors. Before the postsynaptic potentials reach the spike-generation site they are likely low-pass filtered by passive dendritic conduction (Sinz et al., 2020). With the right time-constant this isolates the low-frequency amplitude modulation but does not entirely remove the EOD (Fig. 7, second row). This is supported by the phase locking of P-unit spikes to the EOD. The vector strength quantifying this phase-locking is well below one (Grewe et al., 2017). It would be expected to be much closer to one without low-pass filtering before the spike-generator. With a stronger low-pass filter the P-units would lose their locking to the EOD.

Sensitivity curves measured for a range of EOD frequencies suggest that P-units are tuned to a fish’s EOD frequency (Hopkins, 1976) and to limit P-unit responses to stimulus frequencies close to the EOD frequency. The corresponding band-pass filter is probably caused by electric resonance in the electroreceptor cells (Viancour, 1979). Adding a damped oscillator, Eq. (S27), to our P-unit models, Eqs. (A.1) – (A.4), reproduces P-unit tuning to EOD frequency (Fig. S3 A–B) but does not impair responses to beats at high difference frequencies (Fig. S3 C–D).

### Ambiguity in beat perception

The decline in response amplitude with multiples of the carrier (Fig. 3) results from the declining envelope amplitude (Fig. 6 E). Since the envelope amplitude also depends on the distance between two fish — the larger the distance the smaller the envelope amplitude (Yu et al., 2005) — it can not be used to disambiguate different multiples of *f_EOD_*. Therefore, P-unit responses to similarly mistuned multiples of *f_EOD_* are ambiguous and the fish should not be able to resolve absolute stimulus frequency based on the firing rate modulation of P-units.

The jamming avoidance response (JAR) is evoked by stimulus frequencies close to the receiver’s own EOD frequency and also twice its EOD frequency (Watanabe and Takeda, 1963). Here we report JARs even at the third multiple of the EOD frequency. The EOD frequency shift of the JAR approximately follows the corresponding envelope amplitude as extracted by a cubed threshold up to five times the EOD frequency (Fig. 10). Furthermore, the behavioral threshold for detecting a stimulus frequency (Knudsen, 1974) at least qualitatively follows the repetitive tuning curve of the P-units reported here. This suggests that wave-type electric fish indeed can not disambiguate stimuli at different multiples of their EOD frequency.

In contrast, most fish did not respond to a stimulus frequency close to the zeroth multiple of *f_EOD_*, although there the envelope amplitude is largest. However, also the P-unit response vanished at stimulus frequencies below about 35 Hz (Figs. 1 A,J and 3 C). Such low-frequency stimuli also evoke responses in ampullary cells of the passive electrosensory system (Kalmijn, 1974; Engelmann et al., 2010; Grewe et al., 2017) that would allow the fish to disambiguate the (weak) P-unit responses and to inhibit the JAR response at such low stimulus frequencies.

### Perception of other wave-type species

The wide range of difference frequencies covered by the P-units (Fig. 3) extends far beyond the range of EOD frequencies observed in conspecific wave-type electric fish (usually about one octave). Consequently, the fish should be able to detect the presence of sympatric species that signal in higher or lower frequency ranges (Steinbach, 1970; Hopkins, 1974; Kramer et al., 1981; Stamper et al., 2010; Henninger et al., 2020). Whether and how the different species interact or communicate remains an open question that could be resolved by analyzing electrode-array data recorded in the field (Henninger et al., 2020).

### Beat perception in the auditory system

Early psycho-physical experiments with interacting pure tones demonstrated that beats are not only perceived at low difference frequencies but also for mistuned octaves, when the second tone is close to octaves of the first tone (Roeber, 1834; König, 1876). These experimental findings were formalized in the 19th century by Ohm (1839) and Helmholtz (2009). Beats at higher octaves are better perceived the lower the frequency of the carrier and the louder the signal (Plomp, 1967). Further, masking experiments ruled out aural harmonics and interactions with combination tones as possible mechanisms underlying beat perception (Plomp, 1967). Our observations of clearly modulated firing rate responses at mistuned octaves suggest threshold non-linearities within auditory fibers as a potential mechanism.

### Non-linear physiological mechanisms in the mammalian auditory periphery

Distortion-product otoacoustic emmissions (DPOEs) are hallmark non-linear phenomena of the ear that have been attributed to the mechanical properties of the cochlea and in particular the active amplification of outer hair cells (Brownell, 1990). The most prominent DPOEs are the quadratic distortion, *ω*_2_ − *ω*_1_, which is the difference frequency, and the cubic distortion, 2*ω*_1_ − *ω*_2_ (Kujawa et al., 1995). They could explain beat-like responses to stimuli close to one and two multiples of a tone, but not to higher multiples.

The focus of auditory neuroscience has been on the encoding of SAMs (Joris et al., 2004), often in neurons with quite high characteristic frequencies, such that only the initial declining part of the temporal modulation transfer functions have been recorded (Rhode and Greenberg, 1994). A neuronal correlate of beat perception at high difference frequencies is still not known. Our findings in the electrosensory system predict that a smooth threshold operation within a single auditory nerve fiber could generate distortion products needed to extract the aliasing structure of signal envelopes.

Both the sigmoidal mechanosensory transducer function (Howard and Hudspeth, 1988) as well as the transfer function of the hair-cell ribbon synapse (Moser and Starr, 2016) are good candidates for such a non-linear transformation. In particular, cooperativity of calcium channels in the presynapse has been discussed for hair cells in the auditory system to result in powers of three or higher (Roux et al., 2006; Özcete and Moser, 2020; Peterson and Heil, 2019).

The two superimposed tones need to enter the hair cell with sufficient amplitudes for these mechanisms to take effect. The lower the characteristic frequency of an auditory fiber and the louder the two tones, the wider its effective tuning (Evans, 1972; Sumner and Palmer, 2012), potentially allowing for superimposed tones that differ by multiple octaves to interact within a single auditory fiber. This is in line with the frequency and intensity dependence of beat perception discussed above (Plomp, 1967).

For testing our hypothesis, single auditory fibers should be stimulated with two tones centered symmetrically within their tuning curves. The resulting frequency and amplitude tuning curves allow then to deduce the effective power on the threshold operations implemented in the auditory fiber (Fig. 8). Knock-outs of outer hair cell function or manipulations of the hair cell synapses would then allow to assess the respective contributions.

### Conclusion

A threshold operation with its sharp edge is a mathematical abstraction. Any physiological mechanism implementing this non-linearity, like for example the activation curve of voltage-gated calcium currents or the transfer function of a synapse, has a rather smooth transition. The cubed threshold we derive from our recordings is a mathematically simple way for modeling such a physiologically realistic smooth threshold. In this sense, the ability of the P-units to extract beats at multiples of the carrier frequency is an epiphenomenon of their physiology. For the same reason, mammalian auditory fibers are bound to respond to mistuned octaves and thus should contribute to the percept of beats at higher difference frequencies.

## Appendix A. Methods

## Electrophysiology

40 P-units were recorded from seven weakly electric fish of the species *Apteronotus leptorhynchus* obtained from a commercial tropical fish supplier (Aquarium Glaser GmbH, Rodgau, Germany). The fish were kept in tanks with a water temperature of 25 °C and a conductivity of around 270 μS/cm under a 12 h:12 h light-dark cycle. Body sizes of the fish were between 15 and 17.5 cm and 11.1 and 13.2 g. *f_EOD_* varied between 558 and 860 Hz. All experimental protocols complied with national and European law and were approved by the Ethics Committee of the Regierungspräsidium TÜbingen (permit no: ZP1-16).

## Surgery

Prior to surgery, anesthesia was provided via bath application of a solution of MS222 (120 mg/l, PharmaQ, Fordingbridge, UK) buffered with Sodium Bicarbonate (120 mg/l). For the surgery the fish was fixed on a stage via a metallic rod glued to the skull. The posterior anterior lateral line nerve (pALLN) above the gills, before its descent towards the anterior lateral line ganglion (ALLNG) was disclosed for subsequent P-unit recordings. During the surgery water supply was ensured by a mouthpiece, sustaining anesthesia with a solution of MS222 (100 mg/l) buffered with Sodium Bicarbonate (100 mg/l).

## Experimental setup

Fish were immobilized by an initial intramuscular injection of Tubocurarine (Sigma-Aldrich, Steinheim, Germany; 25–50 μl of 5 mg/ml solution). For the recordings fish were fixated on a stage in a tank, with a major part of the body immersed in water. Analgesia was refreshed in intervals of two hours by cutaneous Lidocaine application (2 %; bela-pharm, Vechta, Germany) around the operation wound and the head mounting rod. Electrodes (borosilicate; 1.5 mm outer diameter; GB150F-8P; Science Products, Hofheim, Germany) were pulled to a resistance of 50–100 MΩ (model P-97; Sutter Instrument, Novato, CA) and filled with 1 M KCl solution. Electrodes were fixed in a micro-drive (Luigs-Neumann, Ratingen, Germany) and lowered into the nerve. Recordings of electroreceptor afferents were amplified (SEC-05, npi-electronics, Tamm, Germany, operated in bridge mode). All signals, P-unit recording, local and global EOD (see below) and the generated stimulus, were digitized with a sampling rate of 40 kHz (PCI-6229, National Instruments, Austin, TX). RELACS (www.relacs.net) running on a Linux computer was used for online spike and EOD detection, stimulus generation, and calibration.

## P-unit identification

P-units were identified based on their firing properties with a baseline firing rate between 64–530 Hz (Grewe et al., 2017; Gussin et al., 2007; Ratnam and Nelson, 2000), phase locking to the EOD, indicated by multimodal interspike-interval (ISI) histograms, and by responses to amplitude modulations of the EOD.

## Electric field recordings

Global EOD for monitoring *f_EOD_* was measured with two vertical carbon rods (11 cm long, 8 mm diameter) in a head-tail configuration. The signal was amplified 200–500 times and band-pass filtered (3 to 1 500 Hz passband, DPA2-FX; npi electronics, Tamm, Germany). A local EOD including the stimulus was measured between two, 1 cm-spaced silver wires located next to the left gill orthogonal to its longitudinal body axis (amplification 200–500 times, band-pass filtered with 3 to 1 500 Hz pass-band, DPA2-FX; npi-electronics, Tamm, Germany). Unfortunately, these filter settings were too narrow for the high stimulus frequencies we used. For Fig. 1 A–F we recreated the local stimulus waveforms by adding the recorded stimulus output to the global EOD to avoid unwanted phase shifts.

## Stimulation

Sine wave stimuli (10–3300 Hz) imitating another fish were isolated (ISO-02V, npi-electronics, Tamm, Germany) and delivered via two horizontal carbon rods located 15 cm laterally to the fish. Depending on *f_EOD_* of the fish, the stimuli resulted in difference frequencies between 750 and 2495 Hz. Each stimulus was repeated twice either for 0.5 s (20% of the trials) or 1 s (80% of the trials). Stimulus amplitude was fixed at 10 % or 20 % of the fish’s local EOD amplitude (contrast) measured prior to each stimulation. From cells stimulated with both amplitudes we only considered the amplitude with the larger number of stimulus frequencies tested for further analysis, such that each cell contributed only once to the population analysis.

## Data analysis

Data analysis was performed with Python 3 using the packages matplotlib, numpy, scipy, sklearn, pandas, nixio (Stoewer et al., 2014), and thunderfish (https://github.com/bendalab/thunderfish).

In binary spike trains with a time step of 0.025 ms each spike was indicated by a value of 40 kHz and all other time bins were set to zero. Time-resolved firing rates were computed by convolving the spike trains with a Gaussian kernel. The standard deviation of the kernels was set to *σ* = 0.5 ms or *σ* = 2 ms. In the frequency domain, these kernels are also Gaussians centered at zero frequency and with a standard deviation of *σ_f_* = (2*πσ*)^−1^ = 318 Hz or *σ_f_* = 80 Hz, respectively. Convolution with kernels corresponds to constructing peri-stimulus time histograms but avoids edge effects introduced by the histogram bins.

Power spectra of binary spike trains or time-resolved firing rates in response were computed from fast Fourier transforms on *n*_fft_ = 4096 long data segments that overlapped by 50 %. The initial and last 5 ms of each spike train were excluded from the analysis.

Frequency tuning curves, the position *f_resp_* of the largest peak in the power spectrum of the time-resolved firing rate as a function of stimulus frequency *f_stim_*, tell us on which frequency the firing rate was modulated by the stimulus. The corresponding amplitude of this peak, *A_resp_*, estimated as the square root of the integral of the power spectrum over the five frequencies closest to the peak frequency, quantifies the modulation depth of this firing rate modulation. Amplitudes of spectral peaks are closely related to the vector strength that is commonly used to quantify temporal modulation transfer functions of auditory neurons in response to SAMs (Sinz et al., 2020). Baseline firing rate was calculated as the number of spikes divided by the duration of the baseline recording (on average 18 s).

During each stimulus presentation we estimated *f_EOD_* from the recorded global EOD as the frequency of the largest peak in power spectra computed with 2^16^ samples per FFT window. In the same way we also confirmed the stimulus frequency. Difference frequencies and stimulus frequencies relative to *f_EOD_* were then reported based on these measurements.

Cells with less than 50 different stimulus frequencies, cells with no stimulus frequencies higher than 2.6*f_EOD_* and cells with no stimulus frequencies between 0 and 100 Hz were excluded from the analysis. Stimulus frequencies resulting in envelopes of periods larger than the analysis window were also excluded from the analysis.

To estimate how far cells follow the aliasing structure to higher stimulus frequencies (*f_max_* in Fig. 3 A), we computed for each stimulus frequency *f_stim_* the quadratic deviation *σ* = (*f_resp_/f_EOD_* − *f_exp_/f_EOD_*)^2^ of the measured response frequency *f_resp_* from the expected aliased frequency *f_exp_* = *f_stim_ f_EOD_ f_stim_/f_EOD_*, where ⌊*x*⌉ rounds *x* to the closest integer. We then looped over all stimulus frequencies *f_stim,i_* and added up the corresponding bins (*f_stim,i_*_+1_ − *f_stim,i_*_−1_)/2 of the stimulus frequencies, but only if the response frequency matched the expected one (*σ*< 0.0005). This sum, *f_max_*, quantifies the range of stimulus frequencies for which the response of the cell follows the expected aliased frequencies.

## Leaky integrate-and-fire models

We constructed leaky integrate-and-fire (LIF) models to reproduce the specific firing properties of P-units (Chacron et al., 2001; Sinz et al., 2020):

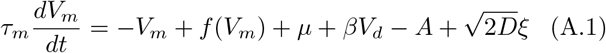

where *τ_m_* is the membrane time constant, *μ* a bias current, and *D* is the strength of Gaussian white noise *ξ*. Whenever the unitless membrane voltage *V_m_* crosses the threshold of *θ* = 1, a spike is generated and the voltage is reset to *V_m_* = 0.

The static non-linearity *f* (*V_m_*) equals zero for the LIF. In case of an exponential integrate-and-fire model (EIF, Fig. 7), this function was set to

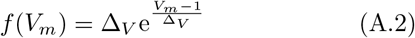

(Fourcaud-Trocmé et al., 2003), where we varied Δ*_V_* from 0.001 to 0.1.

The prominent spike-frequency adaptation of P-units (Benda et al., 2005) is modeled by an adaptation current *A* with dynamics

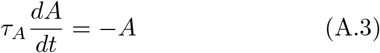

and adaptation time-constant *τ_A_*. Whenever a spike is generated, the adaptation current is incremented by Δ*_A_* (Benda et al., 2010).

The input to the LIF is the membrane voltage *V_d_* of a dendritic compartment scaled by *β*, that low-pass filters the rectified, Eq. (2), electrosensory stimulus *x*(*t*) with a time constant of *τ_d_*:

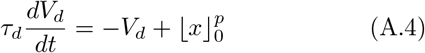

This dendritic low-pass filtering was needed for reproducing the loose coupling of P-unit spikes to the EOD, while maintaining high sensitivity to small amplitude modulations. The rectified stimulus was optionally taken to a power of *p*.

The stimulus is the EOD of the receiving fish normalized to an amplitude of one plus the EOD of a second fish. If not stated otherwise, a superposition of cosine waves, Eq. (1), was used to mimic the EODs. Realistic EODs (Fig. 8 H) were generated by summing up the first 10 harmonics whose relative amplitudes and phases have been extracted from head-tail recordings obtained during measurements of P-unit baseline activity using our thunderfish software, https://github.com/bendalab/thunderfish.

The 8 free parameters of the P-unit model, *τ_m_*, *μ*, *β*, *D*, *τ_A_*, Δ_*A*_, *τ_d_*, and *t_ref_*, were fitted to both baseline activity (baseline firing rate, CV of inter-spike intervals (ISI), serial correlation of ISIs at lag one, and vector strength of spike coupling to EOD) and responses to step in- and decreases in EOD amplitude (onset- and steady-state responses, effective adaptation time constant) of 9 specific P-units (table A.1) for a fixed power of *p* = 1. When modifying the model (e.g. varying the threshold non-linearity or the powers *p*), we adapted the bias current *μ* to restore the original baseline firing rate.

## Jamming-avoidance response

For measuring the jamming avoidance response we placed a fish in a 40 × 50 cm^2^ tank filled with water from their home tank (~250 μS/cm conductivity) to a height of 20 cm, where they voluntarily stayed in a plastic tube. With the same techniques and equipment as for the electrophysiology we measured the fish’s EOD frequency via two carbon electrodes placed in front and behind the tube where the fish was hiding. The fish were stimulated with another pair of carbon electrodes placed orthogonal to the measurement electrodes to the side of the fish about 10 cm apart. Within a time window of 10 seconds the EOD frequency of the fish was estimated from a power spectrum right before each stimulus presentation. Sinewave stimuli were calibrated to an amplitude of 2 mV/cm measured at the position of the fish. Stimulus frequencies were fixed for the 30 sec long duration (no frequency clamping) and set to *k* times *f_EOD_* minus 5 Hz for 1 ≤ *k* ≤ 5 and +5 Hz for *k* = 0, with *f_EOD_* measured right before stimulus onset. Each stimulus frequency was presented once to each fish.

**Table A.1:**
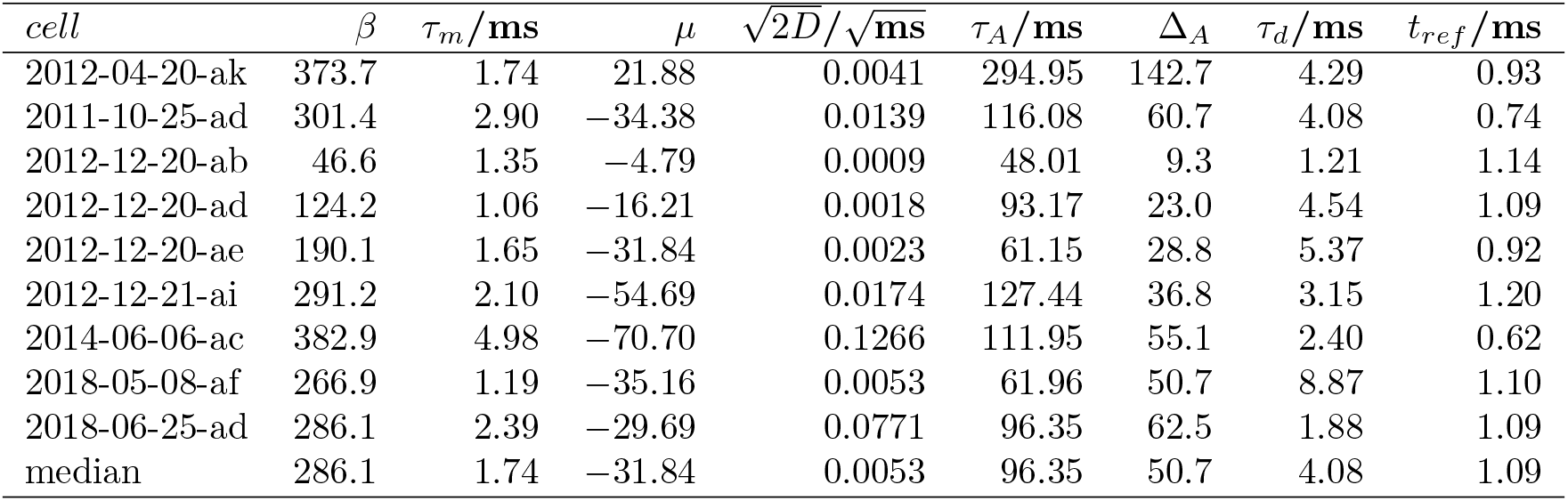
Model parameters, fitted to nine specific P-units.

| *cell* | $\beta$ | $\tau_m/\text{ms}$ | $\mu$ | $\sqrt{2D}/\sqrt{\text{ms}}$ | $\tau_A/\text{ms}$ | $\Delta_A$ | $\tau_d/\text{ms}$ | $t_{ref}/\text{ms}$ |
| --- | --- | --- | --- | --- | --- | --- | --- | --- |
| 2012-04-20-ak | 373.7 | 1.74 | 21.88 | 0.0041 | 294.95 | 142.7 | 4.29 | 0.93 |
| 2011-10-25-ad | 301.4 | 2.90 | -34.38 | 0.0139 | 116.08 | 60.7 | 4.08 | 0.74 |
| 2012-12-20-ab | 46.6 | 1.35 | -4.79 | 0.0009 | 48.01 | 9.3 | 1.21 | 1.14 |
| 2012-12-20-ad | 124.2 | 1.06 | -16.21 | 0.0018 | 93.17 | 23.0 | 4.54 | 1.09 |
| 2012-12-20-ae | 190.1 | 1.65 | -31.84 | 0.0023 | 61.15 | 28.8 | 5.37 | 0.92 |
| 2012-12-21-ai | 291.2 | 2.10 | -54.69 | 0.0174 | 127.44 | 36.8 | 3.15 | 1.20 |
| 2014-06-06-ac | 382.9 | 4.98 | -70.70 | 0.1266 | 111.95 | 55.1 | 2.40 | 0.62 |
| 2018-05-08-af | 266.9 | 1.19 | -35.16 | 0.0053 | 61.96 | 50.7 | 8.87 | 1.10 |
| 2018-06-25-ad | 286.1 | 2.39 | -29.69 | 0.0771 | 96.35 | 62.5 | 1.88 | 1.09 |
| median | 286.1 | 1.74 | -31.84 | 0.0053 | 96.35 | 50.7 | 4.08 | 1.09 |

From the recorded fish’s EOD we computed a spectrogram (FFT segment length of *n* = 131072 samples with an overlap of 50 %). The time course of the EOD frequency was estimated from the peak frequency of the second harmonics in the spectrogram. To compute the frequency shift we subtracted the baseline EOD frequency estimated as the averaged EOD frequency within 10 s right before stimulus onset. Steady-state frequency shift was estimated as the average EOD frequency within 15 to 25 s after stimulus onset relative to baseline EOD frequency.

## Appendix B. Supplement

## 2.1. Analytic signal

The analytic signal corresponding to the original signal is constructed by means of the Hilbert transform. With this method any signal can be expressed as a product

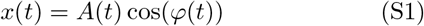

where the amplitude modulation *A*(*t*) is the absolute value of the analytic signal and *φ*(*t*) is the phase of the analytic signal. The amplitude of the carrier cos(*φ*(*t*)) is modulated by *A*(*t*). Whereas the Hilbert transform itself is linear, taking the absolute value is a non-linear operation.

For the superimposed cosines, (1), we get for the amplitude modulation

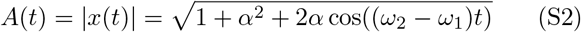

and for the phase

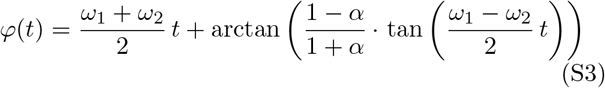

(Stamper et al., 2012). This is an exact identity. The Hilbert transform is just a mathematical trick to transform any signal into such a product of an amplitude modulation and a cosine carrier.

For *α* = 1 (both cosine waves have the same amplitude) this reduces to the well known identity

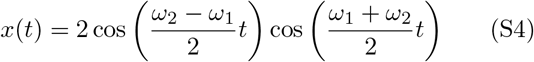

A carrier signal of frequency (*ω*_1_ +*ω*_2_)/2 is multiplied with an amplitude modulation with frequency (*ω*_1_ − *ω*_2_)/2. The latter frequency is half the frequency of the beating amplitude modulation.

For small amplitudes *α* → 0 the expansion of the amplitude modulation to first order results in

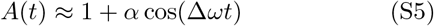

This amplitude modulation has a constant zero-frequency component in the Fourier spectrum, and one at the difference frequency Δ*ω* = *ω*_2_ − *ω*_1_. For larger amplitudes more and more harmonics of this peak appear.

This is exactly what we expect for low difference frequencies, i.e. for stimulus frequencies *ω*_2_ close to *ω*_1_. However, for higher difference frequencies, the amplitude of the analytic signal Eq. (S5) suggests that the beat frequency keeps increasing with increasing difference frequency, no matter how large the difference frequency (Fig. S1 A). It does not explain the aliasing structure we observe in the signals and the P-unit responses. This does not imply that the analytic signal is wrong. Rather the amplitude term (S5) simply does not capture the obvious aliasing structure of the beats. It is hidden in the phase term Eq. (S3).

On a first glance, the phase of the carrier simplifies to *φ*(*t*) = *ω*_1_*t* for small amplitudes. However, this is valid only for *α* = 0, because only then 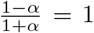 in Eq. (S3). The resulting small-amplitude approximation

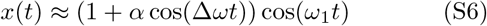

is in fact not a good approximation. In the Fourier spectrum it has two side-peaks at *ω*_1_ ± Δ*ω* instead of only one at *ω*_1_ + Δ*ω* = *ω*_2_ flanking the carrier at *ω*_1_. Eq. (S6) no longer is a beat resulting from the superposition of two cosine waves, but a sinusoidal amplitude modulation (SAM). The approximation fails, because the 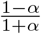-term in Eq. (S3) quickly deviates from one with slope −2 as amplitude increases.

## 2.2. Squaring

An alternative method to retrieve amplitude modulations is to square the signal and then low-pass filter it. Squaring the beat (1), using the binomial theorem and the trigonometric power reduction formula results in

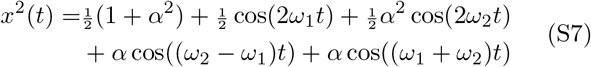

While the original signal (1) has two peaks in the power spectrum at *ω*_1_ and *ω*_2_ and no peak at the beat frequency, the power spectrum of the squared signal (S7) has five peaks, one for each term (Fig. S1 B). Shifting and generating new peaks in the spectrum is a hallmark of non-linear operations. The squaring operation doubles the two original frequencies and creates a new high-frequency peak at the sum of the two frequencies. In addition, a new peak occurs at zero, representing the non-zero mean of the squared signal. Another peak appears at the difference frequency *ω*_2_ − *ω*_1_. This is the amplitude modulation. By subsequent low-pass filtering this peak can be isolated and that way the amplitude modulation can be retrieved. However, as for the analytic signal, none of the five terms explain the aliasing structure of the beat.

## 2.3. Thresholding

The Fourier spectrum of the pulse train, Eq. (4), turns out to have peaks at odd multiples of *ω*_1_ with amplitudes

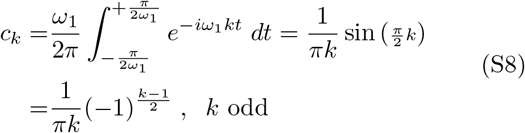

and an additional peak at zero frequency with amplitude

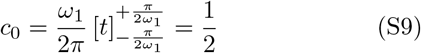

Thresholding a cosine with the same frequency *ω*_1_ can be approximated by multiplying the cosine with the pulse train Eq. (4):

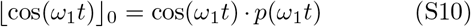

The corresponding Fourier spectrum is the convolution of the spectrum of the cosine with peaks of amplitude 1/2 at ±*ω*_1_ with the spectrum of the pulse train. The two peaks of the cosine are shifted to the positions of all the peaks of the pulse train and multiplied with their amplitude. Always two neighboring peaks of the pulse train at odd multiples of *ω*_1_ contribute to a peak at even multiples of *ω*_1_ with amplitude

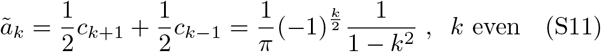

The zero-frequency peak of the pulse train gives rise to peaks at ±*ω*_1_ with amplitude

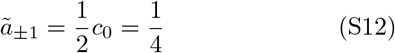

**Figure S1:**
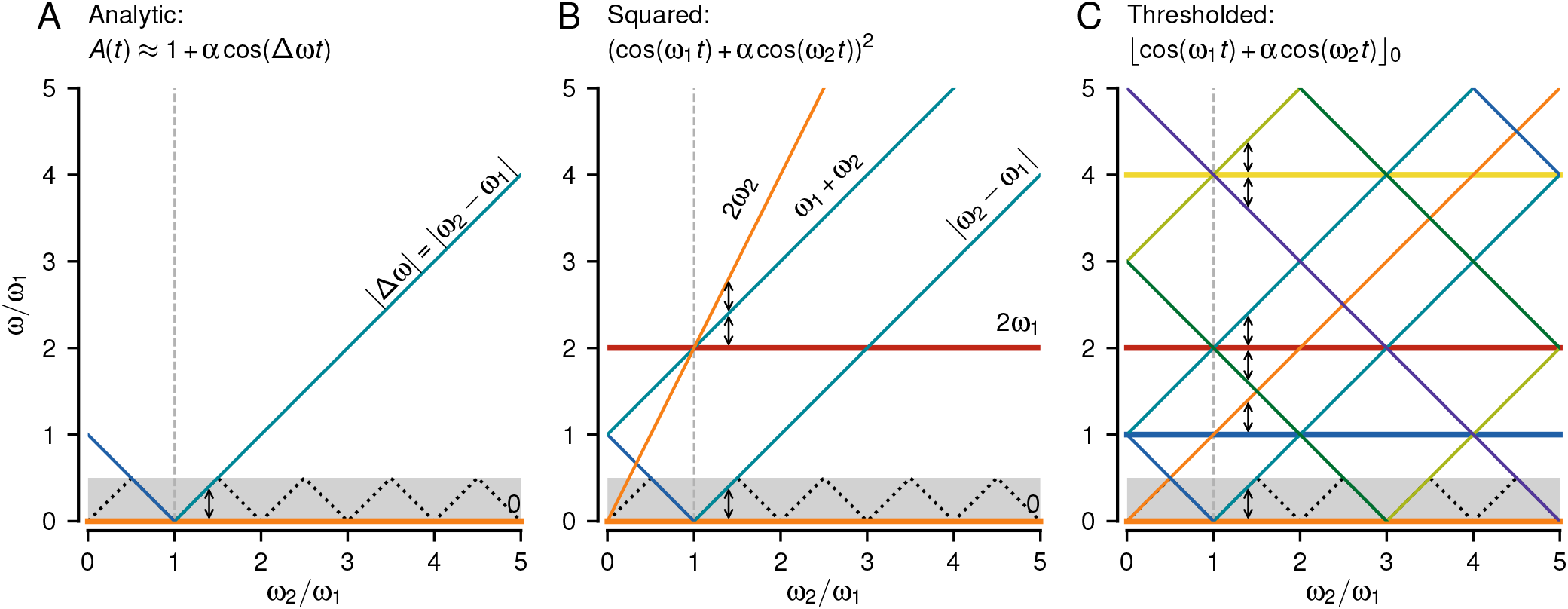
Spectral peaks in analytic, squared and thresholded signals of superimposed cosine waves, Eq. (1). Plotted are the positions of peaks in the spectrum as a function of the stimulus frequency *ω*_2_ relative to the carrier frequency *ω*_1_. For a given stimulus frequency, the corresponding spectrum is a vertical slice through the graph. Vertical arrows highlight the difference frequency Δ*ω* = *ω*_2_ − *ω*_1_. Frequencies below *ω*_1_/2 are marked by the gray background at the bottom. The black dotted line in this frequency band indicates the folded frequencies at *ω_f_* = |*ω*_2_ − *ω*_1_⌊*ω*_2_/*ω*_1_⌉|. **A** The amplitude modulation Eq. (S5) computed as the magnitude of the analytic signal by means of a Hilbert transform has peaks only at 0 and at the absolute difference frequency |Δ*ω*|. **B** Squaring also generates the difference frequency and an offset at zero frequency. In addition, three more peaks appear at 2*ω*_1_, 2*ω*_2_, and *ω*_1_ + *ω*_2_, Eq. (S7). **C** Thresholding the signal results in many more peaks. Convolution of the spectrum of the pulsetrain, Eqs. (S8) and (S9), with the one of the carrier results in horizontal lines at even multiples of *ω*_1_, Eq. (S11), and at *ω*_1_, Eq. (S12). Of interest, however, are the peaks that depend on *ω*_2_. They appear around odd multiples of *ω*_1_, Eq. (S13), and around 0, Eq. (S14).

The spectrum of the thresholded superimposed cosine waves (Fig. S1 C) is composed of the spectrum of the pulse train convolved with the spectrum of the carrier cosine, Eqs. (S11) and (S12), and with the spectrum of the stimulus cosine with peaks of amplitude *α*/2 at frequencies ± *ω*_2_. For the latter, each peak of the pulse train at odd multiples of *ω*_1_ is replaced by a pair of peaks at frequencies *kω*_1_ ± *ω*_2_ with amplitudes

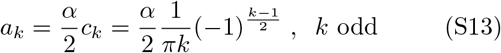

These amplitudes are negative, i.e. they introduce a phase shift by *π*, for every second odd *k* (*k* = 3, 7, 11,…).

The zero-frequency peak of the pulse train, Eq. (S9), gives rise to two peaks at ±*ω*_2_ with amplitude

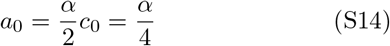

The relative amplitudes *ā_k_* = *a_k_/a*_0_ up to *k* = 5 multiples of *ω*_1_ of the envelope frequencies introduced by thresholding are *ā_0_* = 100 %, 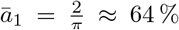, *ā_2_* = 0, 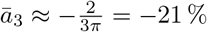, *ā_4_* = 0, and 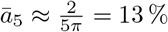.

## 2.4. Threshold cubed

Taking the signal, Eq. (1), to the power of three results in

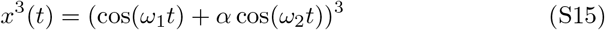

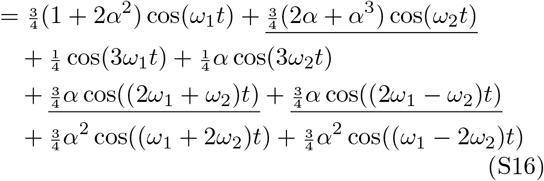

(Fig. S2 A). The dominant peaks depending on *ω*_2_ are at *ω*_2_ and |2*ω*_1_ ± *ω*_2_| (underlined).

Convolving the spectrum of the cubed signal, Eq. (S16), with the one of the pulse train, Eqs. (S8) and (S9), approximating the threshold operation, Eq. (2), boils down to replace all the peaks in the spectrum of the pulse train with the ones of the cubed signal shifted to the respective positions (Fig. 6 A–C). In the following calculations we ignore all terms of higher order in *α*.

The two purely *ω*_1_-dependent terms with peaks at ±*ω*_1_ and ±3*ω*_1_ result in peaks at even multiples of *ω*_1_ with amplitudes

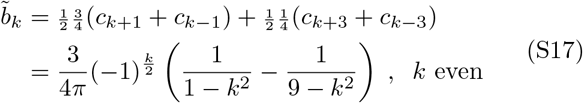

and in addition in peaks directly at ±*ω*_1_ and ±3*ω*_1_ with amplitudes

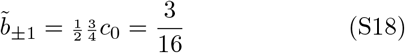

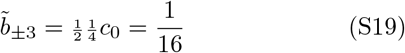

(horizontal lines in Fig. S2 B). The latter at the third harmonics of *ω*_1_ is a new peak that the threshold without exponent does not generate.

**Figure S2:**
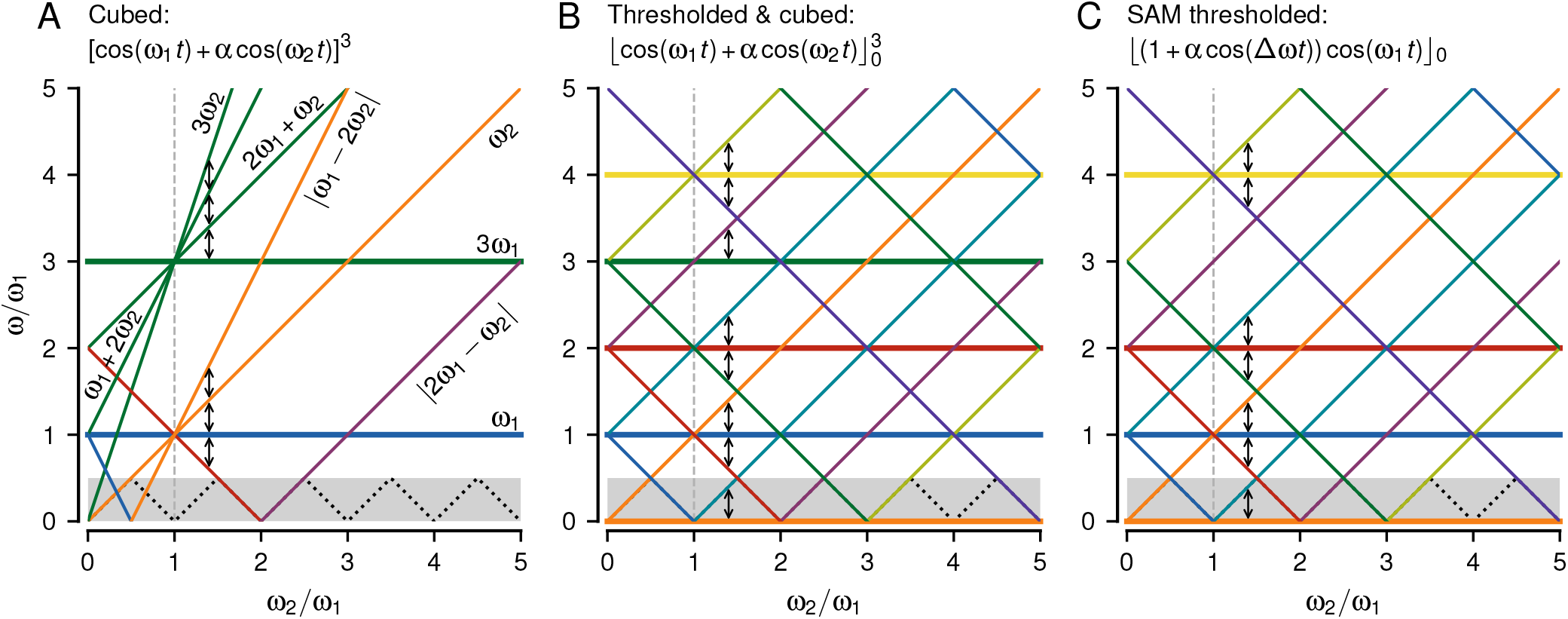
Spectral peaks in cubed and cubed-thresholded superimposed cosine waves as well as for thresholded SAMs. Same style as Fig. S1. **A** Spectral peaks, Eq. (S15), resulting from taking the signal to the power of three. **B** Major peaks resulting from thresholding and cubing the signal. Peaks not depending on *ω*_2_ (horizontal lines) appear at even multiples of *ω*_1_, Eq. (S17), at *ω*_1_, Eq. (S18), and 3*ω*_1_, Eq. (S19). The dominant stimulus-dependent peaks (diagonal lines) appear around odd multiples of *ω*_1_, Eq. (S20), and the zeroth multiple, Eq. (S21), as for the threshold operation without exponent. In addition, however, we get a stimulus-dependent peak around twice the carrier frequency, Eq. (S22). **C** All spectral peaks of a thresholded SAM stimulus, Eq. (S23). Below half the carrier frequency (gray band) it results in the very same peaks as thresholded and cubed superimposed cosines. See supplement section 2.5 for details.

Convolving the dominant *ω*_2_ dependent terms in Eq. (S15) with the peaks at odd multiples of *ω*_1_ of the pulse train, Eq. (S8), we get peaks at *kω*_1_ ± *ω*_2_ for odd *k* with amplitudes

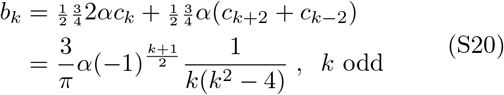

These are peaks at the same frequencies as for the threshold without exponent, but with different amplitudes

However, from the convolution with the zero-frequency term of the pulse train, Eq. (S9), we get additional peaks at ±*ω*_2_ and ±(2*ω*_1_ ± *ω*_2_) with amplitudes

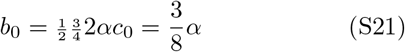

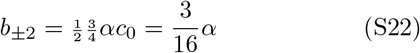

The latter is the one the power of three adds to the folding frequencies around the second multiple of *ω*_1_ (Fig. S2 B).

The relative amplitudes 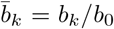 of the envelope frequencies introduced by a cubed threshold are all positive and read 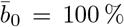, 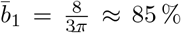, 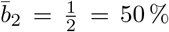, 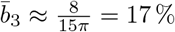, 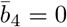, and 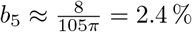.

## 2.5. Thresholding a SAM

A sinusoidal amplitude modulation (SAM) of frequency Δ*ω* and amplitude *α* multiplies a carrier signal with frequency *ω*_1_:

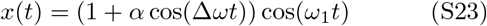

According to the convolution theorem the spectrum of this signal is a convolution of the spectrum of the carrier with peaks at ±*ω*_1_ and amplitude ^1^ with the spectrum of the amplitude modulation with a peak of amplitude 1 at 0 and two peaks at ±Δ*ω* with amplitudes 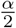. The resulting spectrum of a SAM signal has peaks at ±*ω*_1_ with amplitude 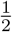, at ±(*ω*_1_ + Δ*ω*) = ±*ω*_2_ with amplitude 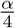, and at ± (*ω*_1_ − Δ*ω*) = ±(2*ω*_1_ − *ω*_2_) also with amplitude 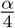. The latter are additional peaks that are not present in the superimposed cosine signal, Eq. (1).

For a SAM the threshold operation, Eq. (2), can be replaced by a multiplication with a pulse train, Eqs. (3) and (4), for all stimulus amplitudes *α*< 1, because the amplitude modulation does not change the zero crossings of the signal. Using the results from above, the convolution of the peaks at ±*ω*_1_ with the pulse spectrum results in peaks at even multiples of *ω*_1_ and at *ω*_1_ with amplitudes Eqs. (S11) and (S12), respectively. The convolution of the peaks at ±*ω*_2_ result in peaks at *kω*_1_ ± *ω*_2_ for odd *k* and *k* = 0 with half of the amplitudes given in Eqs. (S13) and (S14), respectively. The new peaks of the SAM at ±(*ω*_1_ – Δ*ω*) get shifted to the peaks of the pulse train and appear at *kω*_1_ ± (*ω*_1_ − Δ*ω*) = (*k* ± 2)*ω*_1_ ∓ *ω*_2_ for odd *k* with half the amplitudes of Eq. (S13) and for *k* = 0 at ±(2*ω*_1_ – *ω*_2_) with half the amplitude of Eq. (S14). These latter peaks fill in envelope frequencies close to two multiples of *ω*_1_.

## 2.6. Harmonics of the carrier

In reality the carrier EOD is a complex periodic wave and thus already provides harmonics at multiples of the carrier frequency. Wouldn’t that be enough to explain the aliasing structure of the signal envelopes without non-linearities?

At least a threshold is needed. Without any non-linearity the only frequency component depending on the stimulus frequency still would be the stimulus itself. With a threshold, Eq. (2), approximated by multiplication with a pulse train, Eq. (4), the harmonics of the carrier EOD would only add peaks to the resulting spectrum at multiples of the carrier frequency. The stimulus frequency still would be just convolved with the spectrum of the pulse train. As for the sine-wave carrier, the stimulus frequency would appear as envelope frequencies around odd multiples of the carrier frequency and around zero frequency, Eqs. (S13) and (S14), but not at even multiples.

However, the harmonics of the carrier EOD modify the waveform. It is not a sine wave any more that stays positive for exactly half of the time and negative for the other half. Instead, the harmonics might distort the waveform such that we would need a pulse train with a duty cycle other than 50 % to emulate a threshold. For example, some *A. leptorhynchus* have a waveform that is wider than a sine wave at its zero crossings (Fig. 9 A). A matching pulse train would need a higher duty cycle (Fig. 9 B). We parameterize the pulse train by its duty cycle *δ* to account for this effect:

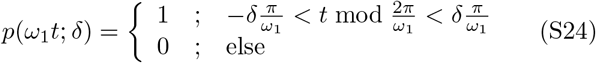

Changing the duty cycle modifies the spectrum of the pulse train:

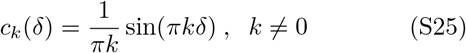

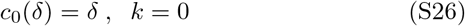

(Fig. 9 B). In particular, a peak at the second multiple of the carrier appears with amplitude 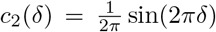.

This peak is then convolved with the stimulus and fills in envelope frequencies around the second multiple (Fig. 9 C). The amplitude of the second multiple of the pulse train equals zero for 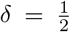, grows linearly in *δ* according to 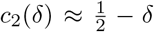 as the duty cycle deviates from 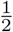 It can get as large as the one of the fundamental, if, according to *c*_2_/*c*_1_ = cos(*πδ*), the duty cycle approaches zero or one. However, the envelope frequencies at the third multiple are missing now, because the third multiple of the pulse train is reduced by increasing the duty cycle.

To summarize, the harmonics of the carrier themselves do not contribute to extracting envelope frequencies. However, the changed duty cycle of the carrier waveform modifies the spectrum of the corresponding pulse train needed to approximate the threshold operation. Depending on the duty cycle of the carrier some harmonics are enhanced whereas others are suppressed.

## 2.7. Harmonics of the stimulus

Alternatively, we could keep the carrier as a sine wave and use a realistic EOD for the stimulus. The components of the signal spectrum relevant for explaining envelope frequencies result from the convolution of the spectrum of a pulse-train with a 50 % duty cycle, Eqs. (S8) and (S9), matching the sinusoidal carrier, with all the harmonics of the stimulus. For extracting the aliasing structure of the envelopes, however, only the fundamental of the stimulus is relevant. The higher harmonics introduce frequencies depending on multiples of the stimulus frequency and thus can not explain the envelope frequencies that grow directly proportionally with stimulus frequency.

## 2.8. Tuning of P-units to EOD frequency

Silencing the fish’s EOD and measuring the minimum amplitude of an artificial replacement EOD to make a P-unit fire action potentials results in V-shaped threshold curves centered at the fish’s EOD frequency (Fig. S3 A, Hopkins, 1976). The corresponding band-pass filter is probably caused by electric resonance in the electroreceptor cells (Viancour, 1979). This could be modeled by a damped harmonic oscillator filtering the input signal before it is thresholded at the receptor synapse (Sinz et al., 2020).

To model this resonance filter we replaced the stimulus *x*(*t*) in the P-unit models, Eqs. (A.1) – (A.4), by the output *y*(*t*) of a harmonic oscillator

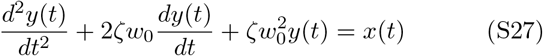

multiplied with a normalization factor *β*. In Eq. (S27) the external force to the oscillator is the stimulus *x*(*t*), *ζ* is the damping ratio of the harmonic oscillator, and

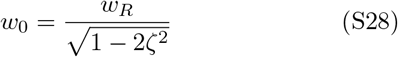

is the eigen-frequency, where *w_R_* = 2*πf_R_* is the resonance frequency that was set to the measured *f_EOD_* of each fish. The normalization factor

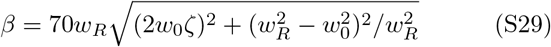

ensures that the fish’s EOD is transmitted with a gain of one through the damped oscillator. The harmonic oscillator was solved using the differential equation solver from SciPy.

**Figure S3:.**
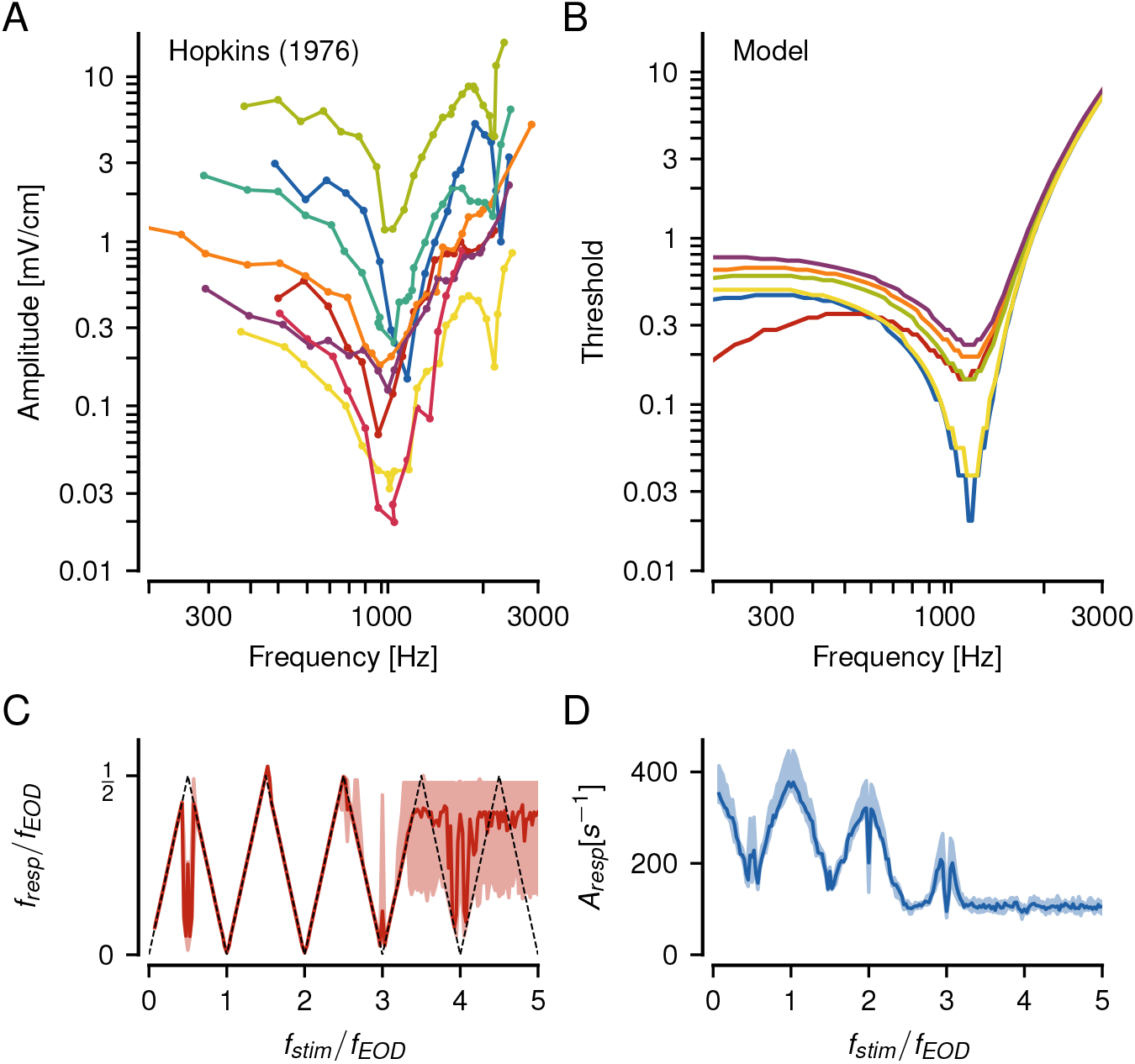
Effects of the P-unit’s EOD frequency filter. **A** Sensitivity of P-unit afferents to EOD frequency as reported by Hopkins (1976). The measured stimulus amplitudes were the minimum amplitude required to elicit just noticeable difference in firing rates of the P-units. **B** Corresponding sensitivities of our LIF models supplemented by a harmonic oscillator Eq. (S27). The amplitudes elicited an increase in firing rate of 10 % compared to baseline rate without stimulus. **C & D** The frequency *f_resp_* and corresponding response amplitudes of these P-unit models to beats still reproduce our observed P-unit tuning to beats despite the EOD filter.

We varied *ζ* from 0.7 (almost no damping) to 0.1 (highest damping). A stronger damping factor of high frequencies can be compensated for by a higher exponent at the threshold, with several combinations yielding similar results.

A mild damping coefficient of *ζ* = 0.45 and an exponent *p* = 5 were sufficient to reproduce both the tuning of P-unit responses to *f_EOD_* as reported by Hopkins (1976) (Fig. S3 B). This model still reproduces the responses to beats up to three multiples of the *f_EOD_* (Fig. S3 C, D), suggesting that the EOD filter does not impede P-unit responses to high difference frequencies.

## Notes

1 Funding: A.B. is funded by DFG grant *BE3699* which is part of the priority program *SPP 2205: “Evolutionary optimization of neural processing“*.

### Competing Interest Statement

The authors have declared no competing interest.

### Summary of Updates

Improved explanation of the phenomenon of slow beat-like envelopes at high difference frequencies. Added new figure 4 and moved figures from the supplement into the main text.

## References

M. S. Lewicki, Efficient coding of natural sounds., Nat Neurosci 5 (2002) 356–363.

C. KÖppl, Phase locking to high frequencies in the auditory nerve and cochlear nucleus magnocellularis of the barn owl, tyto alba, J Neurosci 17 (1997) 3312–3321.

G. L. Romani, S. J. Williamson, L. Kaufman, Tonotopic organization of the human auditory cortex, Science 216 (1982) 1339–1340.

C. D. Hopkins, Stimulus filtering and electroreception: tuberous electroreceptors in three species of gymnotoid fish., J Comp Physiol 111 (1976) 171–207.

A. Roeber, Untersuchungen des Hrn. Scheibler in Crefeld uüber die sogenannten Schl¤ge, Schwebungen oder Stösse, Ann Phys 108 (1834) 492–520.

R. Plomp, Beats of mistuned consonances, J Acoust Soc Am 42 (1967) 462–474.

J. Benda, The physics of electrosensory worlds., in: B. Fritsch, H. Bleckmann (Eds.), The Senses: A Comprehensive Reference, volume 7, Elsevier, Academic Press, 2020, pp. 228–254.

E. Knudsen, Spatial aspects of the electric fields generated by weakly electric fish, J Comp Physiol A 99 (1975) 103–118.

J. Henninger, R. Krahe, F. Sinz, J. Benda, Tracking activity patterns of a multispecies community of gymnotiform weakly electric fish in their neotropical habitat without tagging, J Exp Biol 223 (2020) jeb206342.

N. Yu, G. Hupé, C. Garfinkle, J. E. Lewis, A. Longtin, Coding conspecific identity and motion in the electric sense., PLoS Comput Biol 8 (2005) e1002564.

H. H. Zakon, J. Oestreich, S. Tallarovic, F. Triefenbach, EOD modulations of brown ghost electric fish: JARs, chirps, rises, and dips, J Physiol Paris 96 (2002) 451–458.

G. T. Smith, Evolution and hormonal regulation of sex differences in the electrocommunication behavior of ghost knifefishes (apteronotidae)., J Exp Biol 216 (2013) 2421–2433.

C. E. Carr, L. Maler, E. Sas, Peripheral organization and central projections of the electrosensory nerves in gymnotiform fish., J Comp Neurol 211 (1982) 139–153.

T. Szabo, Sense organs of the lateral line system in some electric fish of the gymnotidae, mormyridae and gymnarchidae, J Morph 117 (1965) 229–249.

A. W. Wachtel, R. B. Szamier, Special cutaneous receptor organs of fish: the tuberous organs of *Eigenmannia*., J Morph 119 (1966) 51–80.

L. Maler, Receptive field organization across multiple electrosensory maps. I. columnar organization and estimation of receptive field size., J Comp Neurol 516 (2009) 376–393.

J. Bastian, Electrolocation I. How electroreceptors of *Apteronotus albifrons* code for moving objects and other electrical stimuli., J Comp Physiol A 144 (1981) 465–479.

M. E. Nelson, Z. Xu, J. R. Payne, Characterization and modeling of p-type electrosensory afferent responses to amplitude modulations in a wave-type electric fish, J Comp Physiol A 181 (1997) 532–544.

J. Benda, A. Longtin, L. Maler, Spike-frequency adaptation separates transient communication signals from background oscillations., J Neurosci 25 (2005) 2312–2321.

P. Joris, C. Schreiner, A. Rees, Neural processing of amplitudemodulated sounds, Physiol Rev 84 (2004) 541–577.

J. Bastian, Electrolocation, J Comp Physiol 144 (1981) 465–479.

J. Benda, A. Longtin, L. Maler, A synchronization-desynchronization code for natural communication signals., Neuron 52 (2006) 347–358.

H. Walz, J. Grewe, J. Benda, Static frequency tuning accounts for changes in neural synchrony evoked by transient communication signals., J Neurophysiol 112 (2014) 752–765.

J. Henninger, R. Krahe, F. Kirschbaum, J. Grewe, J. Benda, Statistics of natural communication signals observed in the wild identify important yet neglected stimulus regimes in weakly electric fish., J. Neurosci. 38 (2018) 5456–5465.

M. J. Chacron, A. Longtin, L. Maler, Negative interspike interval correlations increase the neuronal capacity for encoding time-dependent stimuli, J Neurosci 21 (2001) 5328–5343.

M. Savard, R. Krahe, M. Chacron, Neural heterogeneities influence envelope and temporal coding at the sensory periphery, Neurosci 172 (2011) 270–284.

F. H. Sinz, C. Sachgau, J. Henninger, J. Benda, J. Grewe, Simultaneous spike-time locking to multiple frequencies, J Neurophysiol 123 (2020) 2355–2372.

A. Watanabe, K. Takeda, The change of discharge frequency by A.C. stimulus in a weak electric fish., J Exp Biol 40 (1963) 57–66.

R. König, On the simultaneous sounding of two notes, Phil Mag 1 (1876) 417–446.

G. S. Ohm, Bemerkungen uüber combinationstöone und stösse, Ann Phys 123 (1839) 463–466.

H. L. Helmholtz, On the Sensations of Tone as a Physiological Basis for the Theory of Music, Cambridge University Press, 2009.

H. Scheich, T. H. Bullock, R. Hamstra Jr, Coding properties of two classes of afferent nerve fibers: high-frequency electroreceptors in the electric fish, eigenmannia., J Neurophysiol 36 (1973) 39–60.

N. J. Berman, L. Maler, Inhibition evoked from primary afferents in the electrosensory lateral line lobe of the weakly electric fish *(Apteronotus leptorhynchus)*., J Neurophysiol 80 (1998) 3173–3196.

J. Grewe, A. Kruscha, B. Lindner, J. Benda, Synchronous spikes are necessary but not sufficient for a synchrony code in populations of spiking neurons, P Natl Acad Sci 114 (2017) E1977–E1985.

J. W. Middleton, E. Harvey-Girard, L. Maler, A. Longtin, Envelope gating and noise shaping in populations of noisy neurons., Phys Rev E 75 (2007) 021918.

S. A. Stamper, M. S. Madhav, N. J. Cowan, E. S. Fortune, Beyond the Jamming Avoidance Response: weakly electric fish respond to the envelope of social electrosensory signals., J Exp Biol 215 (2012) 4196–4207.

N. Fourcaud-Trocmé, D. Hansel, C. van Vreeswijk, N. Brunel, How spike generation mechanisms determine the neuronal response to fluctuating inputs, J Neurosci 23 (2003) 11628–11640.

B. W. Knight, Dynamics of encoding in a population of neurons., J Gen Physiol 59 (1972) 734–766.

A. J. Kalmijn, The detection of electric fields from inanimate and animate sources other than electric organs., in: A. Fessard (Ed.), Electroreceptors and other specialized receptors in lower vertrebrates, Springer, Heidelberg, 1974, pp. 148–194.

J. Engelmann, S. Gertz, J. Goulet, A. Schuh, G. von der Emde, Coding of stimuli by ampullary afferents in *Gnathonemus petersii*., J Neurophysiol 104 (2010) 1955–1968.

A. Longtin, J. W. Middleton, J. Cieniak, L. Maler, Neural dynamics of envelope coding, Math Biosci 214 (2008) 87–99.

L. Myers, M. Lowery, M. O’malley, C. Vaughan, C. Heneghan, A. S. C. Gibson, Y. Harley, R. Sreenivasan, Rectification and non-linear pre-processing of emg signals for cortico-muscular analysis, J Neurosci Meth 124 (2003) 157–165.

S. Khanna, M. Teich, Spectral characteristics of the responses of primary auditory-nerve fibers to amplitude-modulated signals, Hearing Res 39 (1989) 143–157.

D.-A. Clevert, T. Unterthiner, S. Hochreiter, Fast and accurate deep network learning by exponential linear units (elus), arXiv preprint arXiv:1511.07289 (2015).

X. Glorot, A. Bordes, Y. Bengio, Deep sparse rectifier neural networks, in: Proceedings of the fourteenth international conference on artificial intelligence and statistics, 2011, pp. 315–323.

A. J. Peterson, P. Heil, Phase locking of auditory-nerve fibers reveals stereotyped distortions and an exponential transfer function with a level-dependent slope, J Neurosci 39 (2019) 4077–4099.

J. W. Middleton, A. Longtin, J. Benda, L. Maler, The cellular basis for parallel neural transmission of a high-frequency stimulus and its low-frequency envelope., P Natl Acad Sci 103 (2006) 14596–14601.

A. Wallach, A. Melanson, A. Longtin, L. Maler, Mixed selectivity coding of sensory and motor social signals in the thalamus of a weakly electric fish., Curr Biol 32 (2022) 51–63.

T. A. Viancour, Electroreceptors of a weakly electric fish, J Comp Physiol 133 (1979) 327–338.

E. Knudsen, Behavioral thresholds to electric signals in high frequency electric fish, J Comp Physiol A 91 (1974) 333–353.

A. B. Steinbach, Diurnal movements and discharge characteristics of electric gymnotid fishes in the Rio Negro, Brazil, Biol Bull 138 (1970) 200–210.

C. D. Hopkins, Electric communication in the reproductive behavior of *Sternopygus macrurus* (gymnotoidei)., Z Naturf 35 (1974) 518–535.

B. Kramer, F. Kirschbaum, H. Markl, Species specificity of electric organ discharges in a sympatric group of gymnotoid fish from Manaus (Amazonas)., Adv Physiol Sci 31 (1981) 195–219.

S. A. Stamper, E. Carrera-G, E. W. Tan, V. Fug`ere, R. Krahe, E. S. Fortune, Species differences in group size and electrosensory interference in weakly electric fishes: Implications for electrosensory processing, Behav Brain Res 207 (2010) 368 – 376.

W. E. Brownell, Outer hair cell electromotility and otoacoustic emissions, Ear Hearing 11 (1990) 82.

S. Kujawa, M. Fallon, R. Bobbin, Time-varying alterations in the f2-f1 dpoae response to continuous primary stimulation i: Response characterization and contribution of the olivocochlear efferents, Hearing Res 85 (1995) 142–154.

W. S. Rhode, S. Greenberg, Encoding of amplitude modulation in the cochlear nucleus of the cat, J Neurophysiol 71 (1994) 1797–1825.

J. Howard, A. J. Hudspeth, Compliance of the hair bundle associated with gating of mechanoelectrical transduction channels in the bullfrog’s saccular hair cell., Neuron 1 (1988) 189–199.

T. Moser, A. Starr, Auditory neuropathy — neural and synaptic mechanisms., Nat Rev Neurol 12 (2016) 189–149.

I. Roux, S. Safieddine, R. Nouvian, M. Grati, M.-C. Simmler, A. Bahloul, I. Perfettini, M. Le Gall, P. Rostaing, G. Hamard, et al., Otoferlin, defective in a human deafness form, is essential for exocytosis at the auditory ribbon synapse, Cell 127 (2006) 277–289.

O. D. Özcete, T. Moser, A sensory cell diversifies its output by varying Ca^2+^ influx-release coupling among active zones., EMBO J 40 (2020) e106010.

E. Evans, The frequency response and other properties of single fibres in the guinea-pig cochlear nerve, J Physiol 226 (1972) 263–287.

C. J. Sumner, A. R. Palmer, Auditory nerve fibre responses in the ferret, Europ J Neurosci 36 (2012) 2428–2439.

D. Gussin, J. Benda, L. Maler, Limits of linear rate coding of dynamic stimuli by electroreceptor afferents, J Neurophysiol 97 (2007) 2917–2929.

R. Ratnam, M. E. Nelson, Nonrenewal statistics of electrosensory afferent spike trains: implications for the detection of weak sensory signals, J Neurosci 20 (2000) 6672–6683.

A. Stoewer, C. J. Kellner, J. Benda, T. Wachtler, J. Grewe, File format and library for neuroscience data and metadata, Front Neuroinform 8 (2014) 10–3389.

J. Benda, L. Maler, A. Longtin, Linear versus nonlinear signal transmission in neuron models with adaptation-currents or dynamic thresholds., J Neurophysiol 104 (2010) 2806–2820.

